# The transcriptome of regenerating zebrafish scales identifies genes involved in human bone disease

**DOI:** 10.1101/2020.10.08.331561

**Authors:** Dylan J.M. Bergen, Qiao Tong, Ankit Shukla, Elis Newman, Jan Zethof, Mischa Lundberg, Rebecca Ryan, Scott E. Youlten, Eleftheria Zeggini, Peter I. Croucher, Gert Flik, Rebecca J. Richardson, John P. Kemp, Chrissy L. Hammond, Juriaan R. Metz

## Abstract

Zebrafish scales are mineralised plates that can regenerate involving *de novo* bone formation. This presents an opportunity to uncover genes and pathways relevant to human musculoskeletal disease relevant to impaired bone formation. To investigate this hypothesis, we defined transcriptomic profiles of ontogenetic and regenerating scales, and identified 604 differentially expressed genes (DEGs) that were enriched for extracellular matrix, ossification, and cell adhesion pathways. Next, we showed that human orthologues of DEGs were 2.8 times more likely to cause human monogenic skeletal diseases (P<8×10^−11^), and they showed enrichment for human orthologues associated with polygenetic disease traits including stature, bone density and osteoarthritis (P<0.005). Finally, zebrafish mutants of two human orthologues that were robustly associated with height and osteoarthritis (*COL11A2*) or bone density only (*SPP1*) developed skeletal abnormalities consistent with our genetic association studies. *Col11a2*^*Y228X/Y228X*^ mutants showed endoskeletal features consistent with abnormal growth and osteoarthritis, whereas *spp1*^*P160X/P160X*^ mutants had elevated bone density (P<0.05). In summary, we show that transcriptomic studies of regenerating zebrafish scales have potential to identify new genes and pathways relevant to human skeletal disease.

## Introduction

The zebrafish skeleton, similar to higher vertebrates, provides structure, organ protection, endocrine and locomotive function. During development, bone is formed by two evolutionarily conserved processes via either endochondral or intramembranous (dermal) ossification. Endochondral bone formation occurs though progressive remodelling of a cartilage template into bone, while dermal or intramembranous bone is formed directly by mesenchymal condensation of osteoblasts. Due to mechanical loading induced microfractures, bone is a dynamic tissue that undergoes constant remodelling of the calcified extracellular matrix (ECM) via a tight coupling between the functional activity of the bone-building osteoblasts and bone-degrading osteoclasts, allowing a regenerative capacity of bone throughout life (Kenkre & Bassett, 2018). In common age-related conditions such as osteoporosis, this balance is dysregulated such that catabolism exceeds anabolism, making bone more fragile and susceptible to fracture. Current pharmacotherapies for osteoporosis are limited, with most targeting the catabolic activity of osteoclasts rather than on bone anabolism. However, human genetic studies of rare bone diseases have shed light on potential bone anabolic pathways leading to development of novel therapeutics (Pathak, Bravenboer et al., 2020). For example, linkage studies have identified the WNT pathway inhibitor Sclerostin (SOST) that leads to Sclerosteosis causing high bone mass (HBM) in humans and mice when mutated (Balemans, Ebeling et al., 2001, Li, Ominsky et al., 2008). The subsequent development of a humanised monoclonal antibody (Romozumab) against SOST now offers an osteo-anabolic therapeutic that reduces fracture risk for osteoporosis patients (Cosman, Crittenden et al., 2016). This demonstrates how understanding the genetic control of bone growth and osteoblast differentiation processes allow identification of novel therapeutic targets.

One potential route to identifying new pathways and targets is through the study of organisms and tissues capable of rapid regrowth of skeletal tissues, during which bone anabolism exceeds bone catabolism. While skeletal regenerative capacity in mammals is limited to bone remodelling and fracture healing, a number of other vertebrates species such as zebrafish are able to undergo epimorphic regeneration of many organs throughout life, including the bony structures of the fins and scales (Zhao, Qin et al., 2016). Zebrafish are increasingly used as a model for musculoskeletal (MSK) research, due to their genetic tractability, availability of transgenic reporter lines and the dynamic imaging opportunities, they offer as well as the similarities of their skeletal physiology to humans and other higher vertebrates (Lleras-Forero, Winkler et al., 2020). Zebrafish fin regeneration has been extensively studied and has revealed many of the mechanisms that underpin blastema formation and skeletal dedifferentiation and re-differentiation during tissue regrowth (Sehring & Weidinger, 2020).

Most teleost fishes have scales which function as a protective armour and a calcium reservoir (Sire, Donoghue et al., 2009, Yasuo, 1980).The zebrafish elasmoid scales, that also undergo complete epimorphic regeneration within days to weeks, have recently gained attention in studies aiming at deeper understanding of mechanisms of bone growth, regeneration, and repair. In common with intramembranous flat bones, the elasmoid scales are formed and mineralised directly by *de novo* differentiated osteoblasts (Pasqualetti, Banfi et al., 2012, Sire, Allizard et al., 1997). Scales are exoskeletal elements that have been strongly reduced or completely lost in terrestrial animals during evolution but have been retained in bony fish. Long regarded to be odontogenic in origin, relatively recent studies have provided new insights into the classification of skeletal structures and evolutionary and developmental origins of exoskeletal teleost scales (Dhouailly, Godefroit et al., 2019, Shimada, Kawanishi et al., 2013). Following removal of the ontogenetic scale from the dermal socket initiating a wound healing inflammation phase, regeneration is initiated and a small mineralised scale plate can be observed as early as two days post-harvest (dph) (Bereiter-Hahn & Zylberberg, 1993, Richardson, Slanchev et al., 2013, Sire & Akimenko, 2004). Regenerating scales have a high density of osteoblasts at the posterior edge of the plate forming the leading growth plane whose dynamics can be tracked *in toto* (Cox, De Simone et al., 2018). These osteoblasts form a (hyposquamal) monolayer and deposit hydroxyapatite into a type I collagen rich matrix that can be resorbed by osteoclasts (de Vrieze, Sharif et al., 2011, Guellec & Zylberberg, 1998). The collagen matrix of scales is organised in a plywood manner resembling lamellar bone in humans (Bergen, Kague et al., 2019, Bigi, Burghammer et al., 2001, Giraud-Guille, 1988). Similar to human bones, scale bone turnover responds to exposure of prednisolone, alendronate, chronic hyperglycaemia, and fatty acids (e.g. OMEGA-6) highlighting evolutionarily conserved mechanisms of bone metabolism (Carnovali, Luzi et al., 2016, Carnovali, Ottria et al., 2016, de Vrieze, van Kessel et al., 2014, Pasqualetti, Congiu et al., 2015). Moreover, ontogenetic scales show a fracture healing response with recruitment of *trapc* expressing cells to the fracture site (Kobayashi-Sun, Yamamori et al., 2020). The possibility to *ex vivo* culture fish scales as small transparent bone units, in which vital intercellular interactions (between osteoblasts and osteoclasts), as well as cell-matrix interactions remain intact, has triggered their use in screenings for osteogenic compounds (de Vrieze et al., 2011, de Vrieze, Zethof et al., 2015). Hence, these features therefore warrant a deeper analysis of genes, pathways in the scale and a comparison with genes associated with human MSK disease.

As regeneration leads to rapid bone regrowth we hypothesised that (i) osteoanabolic genes can be identified by contrasting the transcriptomic profiles of ontogenetic and regenerating zebrafish scales, and (ii) that these identified “bone formation” genes were likely to be enriched for human orthologues that influence bone growth, mineralisation and susceptibility to MSK disease. To test our hypotheses, we identified differentially expressed genes (DEGs) that distinguished regenerating scales from ontogenetic scales, identified biological pathways that DEGs were involved in, and investigated whether DEGs were associated with human monogenic disease, and polygenetic disease traits: height, bone mineral density and osteoarthritis susceptability. Lastly, we performed functional studies of two DEGs that were robustly associated with one or more human MSK disease traits and observed abnormal endoskeletal phenotypes consistent with the human genetic association analyses.

## Results

### Defininig the transcriptome of ontogentic and regenerating scales by RNA-sequencing

We first confirmed that,regenerating scales show an increase in the number of *sp7* positive osteoblasts compared to original scales formed during development (ontogenetic) as previously reported (Cox et al., 2018, de Vrieze et al., 2014) (**figure 1A**). Alkaline phosphatase (ALP) staining showed that regenerating scales contained more ALP activity (**figure 1B**). To identify the biological pathways underpinning the regeneration process, we preformed RNA-sequencing (RNA-seq) on ontogenetic (original) and regenerating scales 9 days into their regeneration (**figure 1C**).

**Figure 1:**
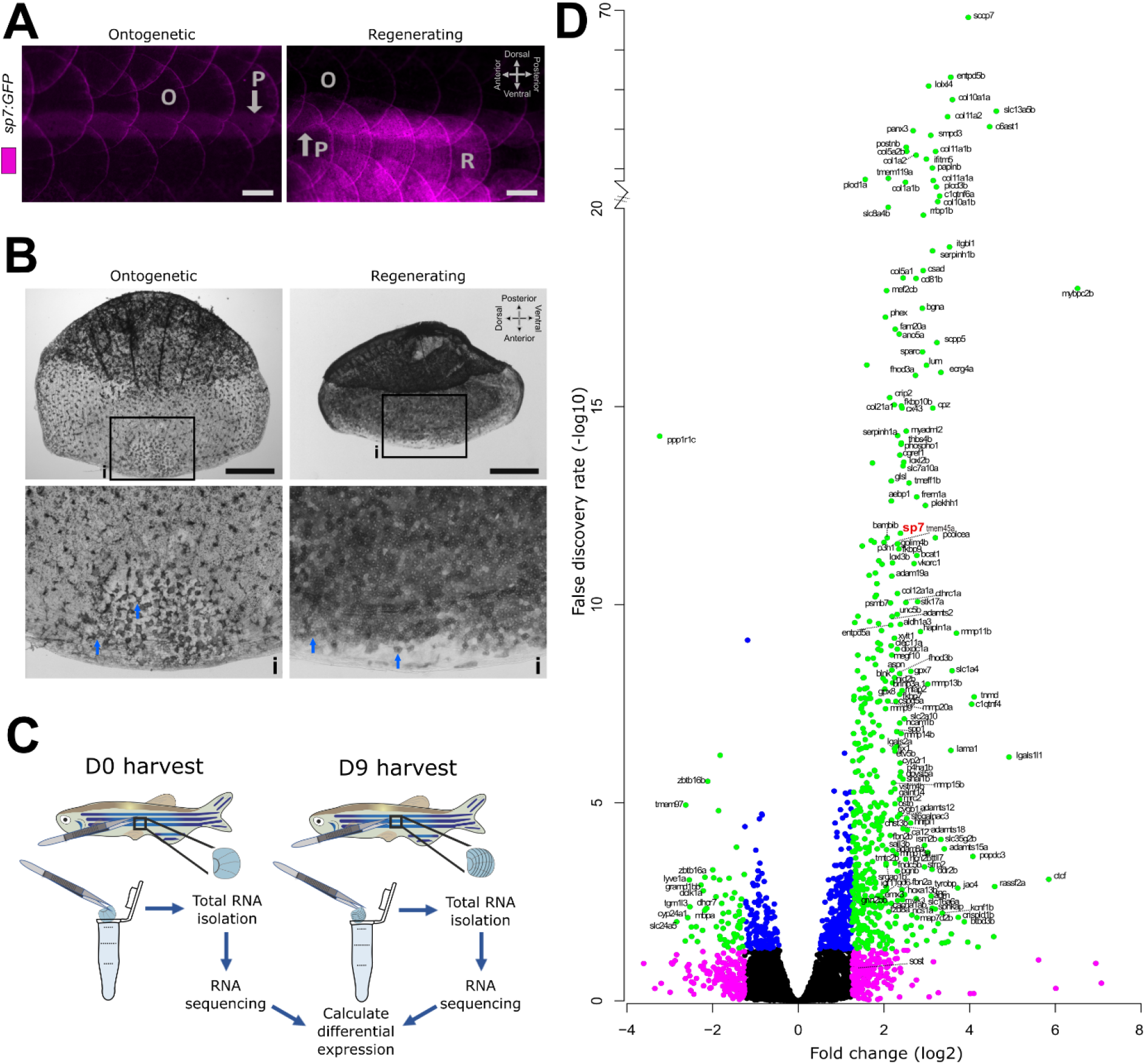
RNA-sequencing transcriptomics of ontogenetic and 9 days regenerated scales. **A)** *In toto* fluorescent stereomicroscope images of flanks from *sp7:GFP* transgenic fish showing ontogenetic (O) and regenerated scales (R) in an anterior (left) to posterior (right) direction). Note the auto-fluorescent pigment stripe (P) and the compass orientation is used to depict *in toto* scales in this paper with the distal (posterior) edge pointing caudally. **B)** *In situ* alkaline phosphatase staining of a pre-collection and 8 dph scale with insets (i) showing anterior part of the scale. Blue arrows indicate ALP+ cells which appear to be larger and more extended in ontogenetic than regenerating scales. Compass orientation is used to depict *in situ* scales in this paper as the anterior region was closest to the scale dermal socket (pre-harvest). **C)** Schematic drawing of the RNA-sequencing approach. **D)** Volcano plot showing relative fold change (log2) and false discovery rate (-log10 converted) of coding RNA sequences expressed in both ontogenetic and regenerating scales. Green (≥±1.25 fold change, ≥ 1.3 FDR), magenta (≥±1.25 fold change, <1.3 FDR), blue (<±1.25 fold change, ≥1.3 FDR), and black (not passing any threshold) coloured dots mark the different criteria.

A total of 13,170 protein coding genes were consistently expressed (i.e. 51.4% of total protein coding genome) in both ontogentic and regenerating scales (**data file S1**). Principal component analysis (PCA) of the two groups of all genes expressed showed that ontogenetic and regenerating scale groups cluster together (**figure S1A**) and that there was a a high level of correlation (Pearson’s correlation of 97.1%, **figure S1B**), confirming similar expression patterns within the PCA groups. Setting the arbitrary threshold at 1.25 log2 fold change and a false discovery rate (FDR) of <0.05 showed that 604 protein coding genes had substantial differences in gene expression (**data file S2**). We observed a skew towards upregulation of genes (*n = 514* compared to *n = 90* downregulated); consistent with regenerating tissues such as the caudal fin (Padhi, Joly et al., 2004, Schmidt, Geurtzen et al., 2019) (**figure 1D**). The osteoblast transcription factor *sp7(osxterix)* was substantially upregulated (2.39 log2 fold change) in regenerating scales (**figure 1D**), but 128 genes showed a larger up regulation than *sp7*. Among the top DEGs, we noticed a high number of collagen and cell adhesion related genes (**table 1**).Two genes were highly downregulated and a homology search revealed that both genes encoded different proteins [i.e. ENSDARG00000068621 (*si:ch211-181d7.3*) and ENSDARG00000088274 (*si:ch211-181d7.1*)] that were unique to fish(**figure S2**). Both genes have NOD-like receptor (NLR) domains that possess NACHT (NTPase) and leucine-rich repeat (LRR) protein domains important for innate immune system antigen interactions (Wu, Chen et al., 2018).

**Table 1:**
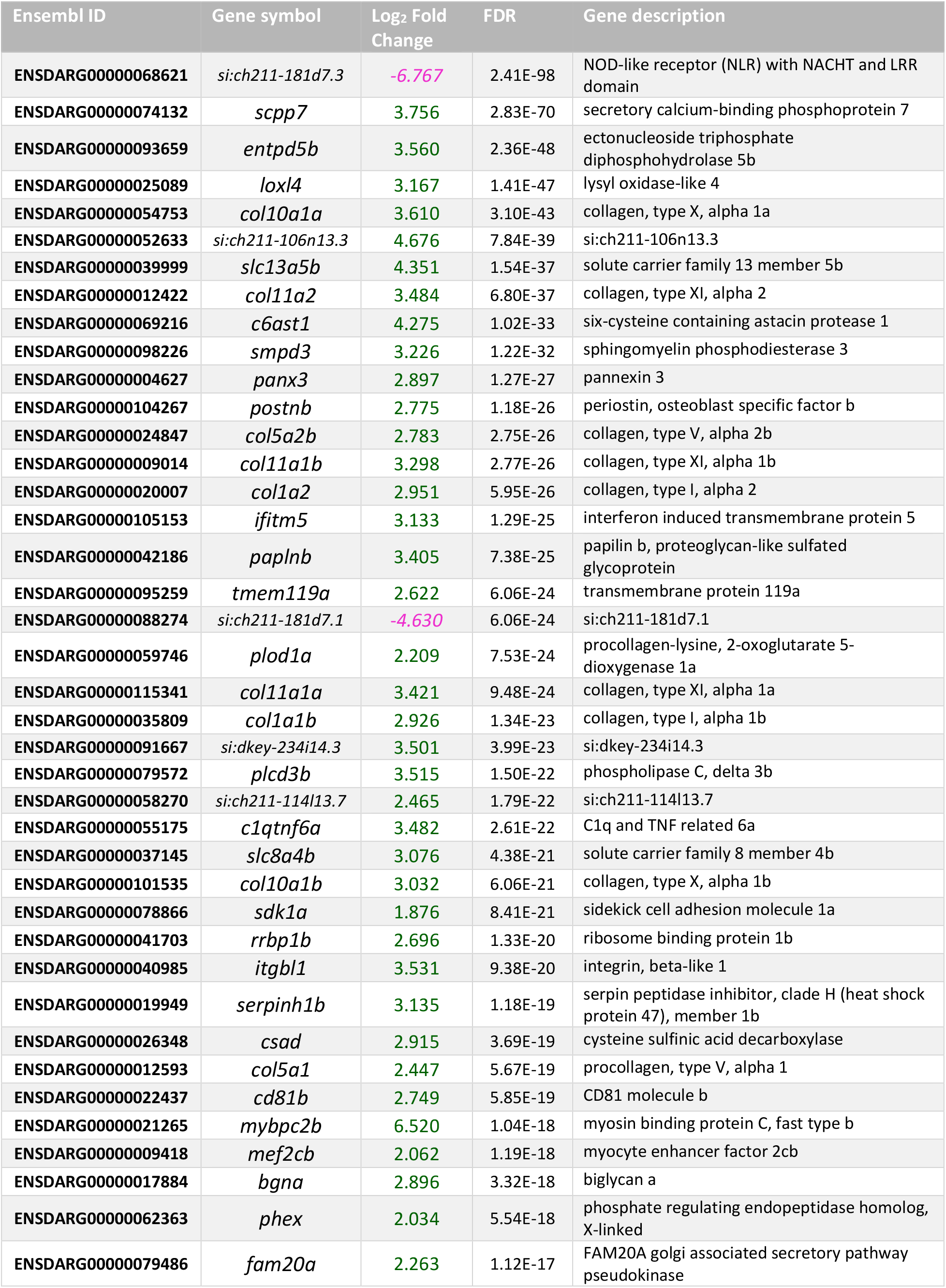
Top-40 smallest false discovery rate (FDR) differentially regulated transcripts (up (green) and down (magenta) regulated in Log2 fold change) in regenerated scales (vs ontogenetic) showing mean and standard deviation (Stdev) for each group (normalised read count).

Osteoblast factors such as *sp7, entpd5a/b, postnb, smpd3, phospho1, plod1a, plod2,* and *plod3* were upregulated in 9 dph scales, as were genes related to ECM growth, collagens (predominantly fibrillar collagens) and collagen remodelling including *bmp1a*, *mmp9*, *spp1*, *ostn*, *col1a1a/b*, *col1a2*, *col5a1, col5a2a/b, col10a1a/b*, *bgna/b*, *col11a1a/b* and *col11a2* (Ricard-Blum, 2011). We also observed elevated expression of genes normally associated with endochondral ossification: *col11a1a/b* and *col11a2, ihha, ptch2, dlx5a,* and *mef2cb*, suggesting that some genes are common to both modes of ossification (Long & Ornitz, 2013). While osteoclast and monocyte related factors (e.g. *tnfrsf11a* (*RANK*), chloride channel *clc7n7, cathepsin-K* (*ctsk*), and *acp5a/b (trapc*)) were expressed in both ontogenetic and regenerating scales, differential expression was not observed (**data file S1**). To validate the RNA-seq dataset, quantitative real-time PCR (qRT-PCR) was performed on osteoblast and osteoclast markers and showed concordant data between RNA-seq and qRT-PCR assays of all selected amplicons (**figure 2A**). These results indicate osteoblast activity is elevated to a greater degree than osteoclast activity. To visualise newly formed bone, *in vivo* calcein green staining was performed (**figure 2B**). The calcein green signal was stronger in regenerating scales, and predominantly seen in the posterior regions of the scale, consistent with elevated expression of bone anabolic factors such as *sp7* and *entpd5a* (**figure 2A** and **2B**).

**Figure 2:**
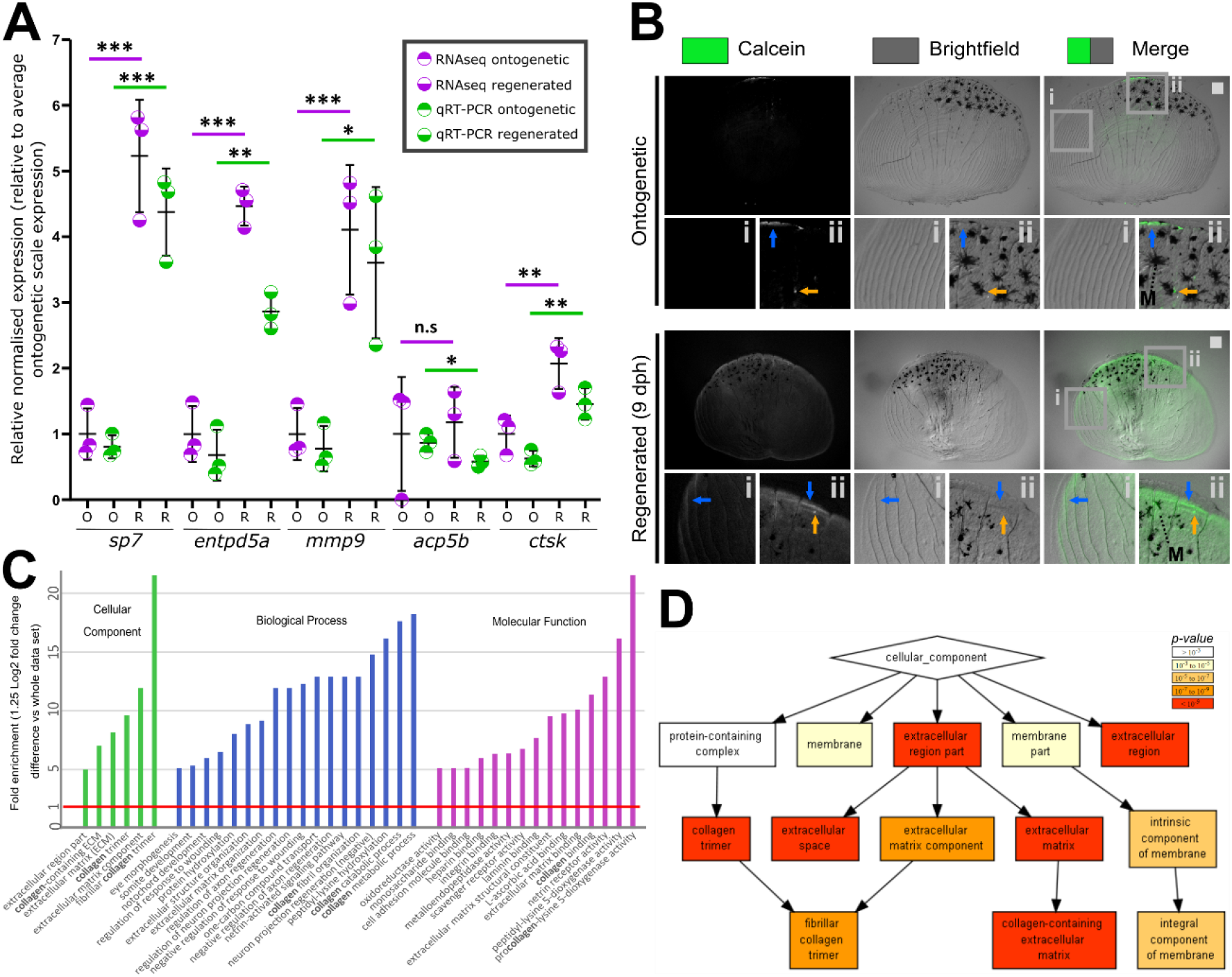
Validation and gene ontology of RNA-sequencing dataset. **A)** Comparison of relative expression levels of quantitative real-time PCR (qRT-PCR) and transcriptomic analysis of selected amplicons. **B)** Representative stereomicroscope images of live calcein green staining labelling newly deposited calcium phosphate of ontogenetic and regenerating scales (n = 4 fish each condition). Blue arrow indicates elevated levels of calcein green labelling compared to surrounding signal whereas orange arrows indicate small puncta of enhanced signal. Insets show lateral circuli (i) and posterior epidermal (containing pigmentating melanocytes (M) region (ii). **C)** GOrilla Gene ontology (GO) analysis showing high enrichment (>5 fold) of collagen and ECM related terms. **D)** GOrilla cellular component hierarchical clustering. Scale bar: 100 μm.

### The collagen processing pathway is highly enriched and upregulated during scale regeneration

To identify biological pathways overrepresented among DEGs, we performed gene ontology (GO) enrichment analysis. Several, bone specific GO terms were significantly overrepresented among DEGS, such as ‘regulation of bone biomineralization’ (GO:0110149; 10.9-fold, FDR=0.018), ‘ossification’ (GO:0001503; 6.0-fold, FDR=0.0019), and ‘skeletal system development’ (GO:0001501; 3.6-fold, FDR=5.3×10^−6^). Collagen related terms were also significantly enriched (>7.1 fold, FDR=8.7×10^−6^) (**figure 2C, data file S3**), including ‘collagen trimer’, ‘collagen-containing extracellular matrix’ and ‘fibrillar collagen trimer’ GO terms (**figure 2D**). Additional bone matrix associated processes such as ‘calcium-ion binding’, ‘glycosaminoglycan binding’ and. ‘matrix metalloprotease (MMP) activity’ were identified (**figure S3 and S4**). Further, the ‘integrin signalling pathway’ (P00034; 3.5-fold, FDR=0.016) was the only enriched signalling pathway, which is known to bind collagen during cell adhesion related processes (Zeltz & Gullberg, 2016) (**data file S4**). Together, this indicated DEGs were enriched for biological pathways involved in bone formation and collagen-matrix synthesis, consistent with increased osteoblast activity.

We next visualised protein-protein interactions (PPI) within this set using STRING Network analysis. This broadly confirmed GO findings and showed that the DEGs PPI network has 6.6-fold higher number of nodes (connections) compared to the whole reactome (P<1.0×10^−16^). There were ten clusters with distinct functions that were significantly enriched that were highly associated with collagen-rich ECM and cell adhesion (**figure 3**). Additional terms not identified by GO included ‘regulation of insulin-like growth factor (IGF) signalling’ and ‘signalling by hedgehog’ (HH) (**figure 3**). Note that STRING added 10 ‘high-scoring’ interactors to the network that were not DEGs (**figure S5A**). There were four clusters unlinked to the main network, related to cytoskeletal structure or cell motility such as actin-myosin (cluster 1 and 4), cell motility proteins that regulate leading edge cytoskeletal modelling (cluster 2), and enzyme function (cluster 3) (**figure 3** and **table S1**). Allowing less stringent interaction evidence scores (≥ 0.4) showed that *sp7* and *secreted phospho protein 1* (*spp1*, encoding Osteopontin (Opn)) factors have connections with both the collagen and bone factor clusters, the latter showing many connections to adjacent osteogenic clusters that for example contain *phospo1* and *ostn* (**figure S5B**). These findings were verified with PANTHER ‘Reactome pathways’ analysis (**data file S5**). These GO and PPI analyses showed that the RNA-seq dataset is specific for an osteogenic collagen rich regenerating tissue.

**Figure 3:**
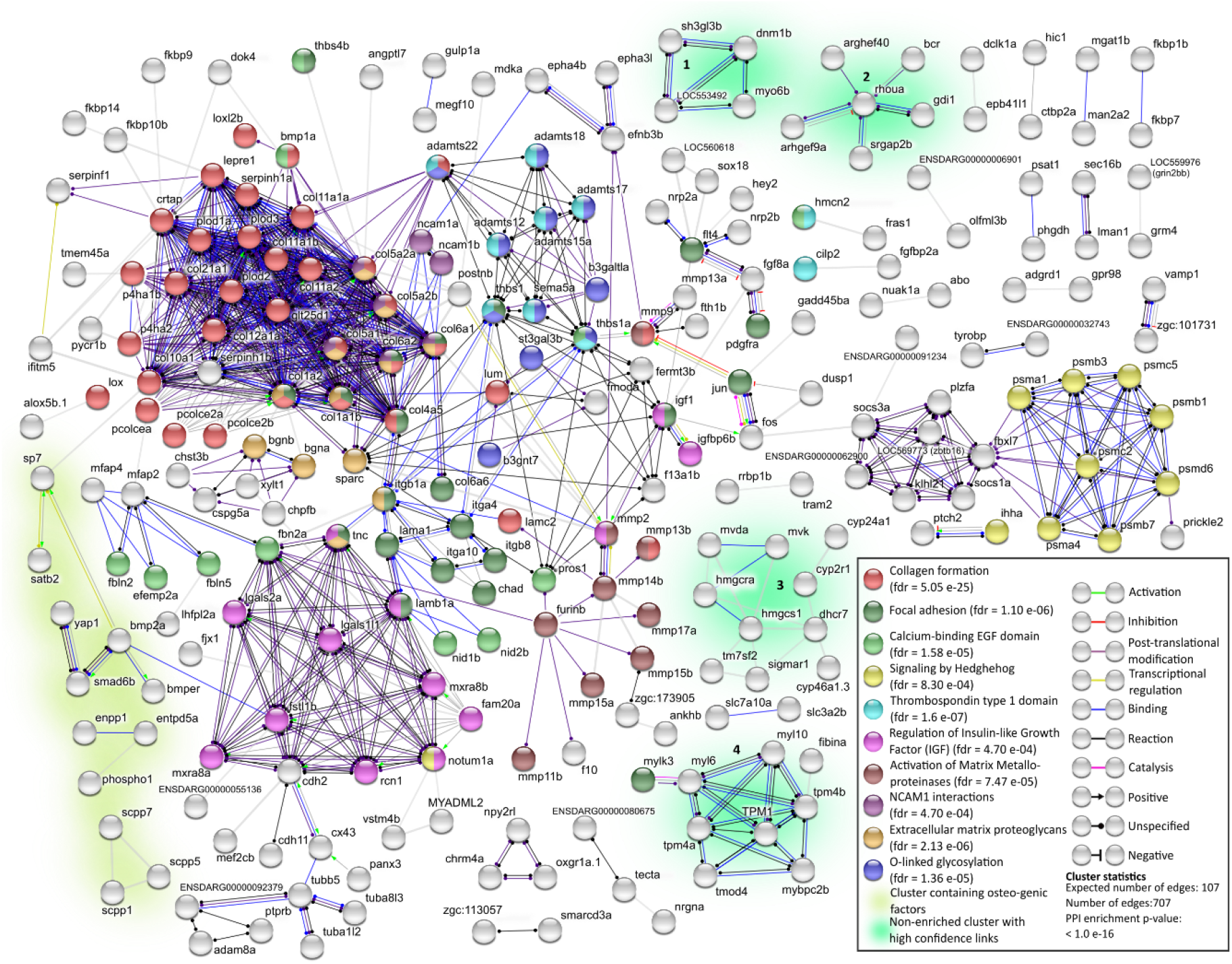
STRING network analysis of DEGs. Only DEG proteins with 1 or more protein-protein interactions (PPI) (indicated as edges) were shown.

We next verified RNA expression of *col1a2,* osteoblast deposited collagen *col10a1a,* the collagen nucleation factor proteoglycan *bgna,* and hedgehog signalling ligand *ihha* that represent some of these clusters identified above. The expression of the gap junction gene *cx43* was also assessed as it regulates fin regeneration growth, and within our PPI network it was connected to the IGF cluster known to modulate bone growth (Bhattacharya, Hyland et al., 2020). These amplicons were assessed in regenerating scales from an independent experiment and all showed similar trends between RNA-seq and qRT-PCR, validating these findings (**figure 4A**). Note, a subset of genes showed similar profiles between the RNA-seq and independent experiments (**figure S6**). Ontogenetic scales showed weak *col1a1a* expression, predominantly located in the epidermis adjacent to *sp7* positive cells at the posterior edge of the scale (**figure 4B**). In regenerating scales, *col1a1a* promoter activity was elevated along with expanded *sp7* expression covering most of the bony unit. We observed elevated activity at newly forming circuli associated with thickening of the calcified ECM (**figure 2B** and **figure 4B**). We performed the same procedure in *col10a1a:Citrine* and *col2a1a:mChrerry* double transgenic zebrafish where the *col2a1a* reporter functions as a negative control as it was not a DEG. This showed that ontogenetic reporter expression of both transgenes was absent. At 9 dph, *col10a1a* reporter expression was observed at the posterior edge of the scale and interestingly also at the newly developing circuli (thicker ECM) located at the anterior region of the scale (**figure 4C**). A mCherry signal was not detected in ontogenetic or regenerating scales. These findings confirm that the RNA-seq dataset of regenerating scales contains gene networks with osteo-active factors that actively assemble (nucleate) a calcified collagen-rich matrix.

**Figure 4:**
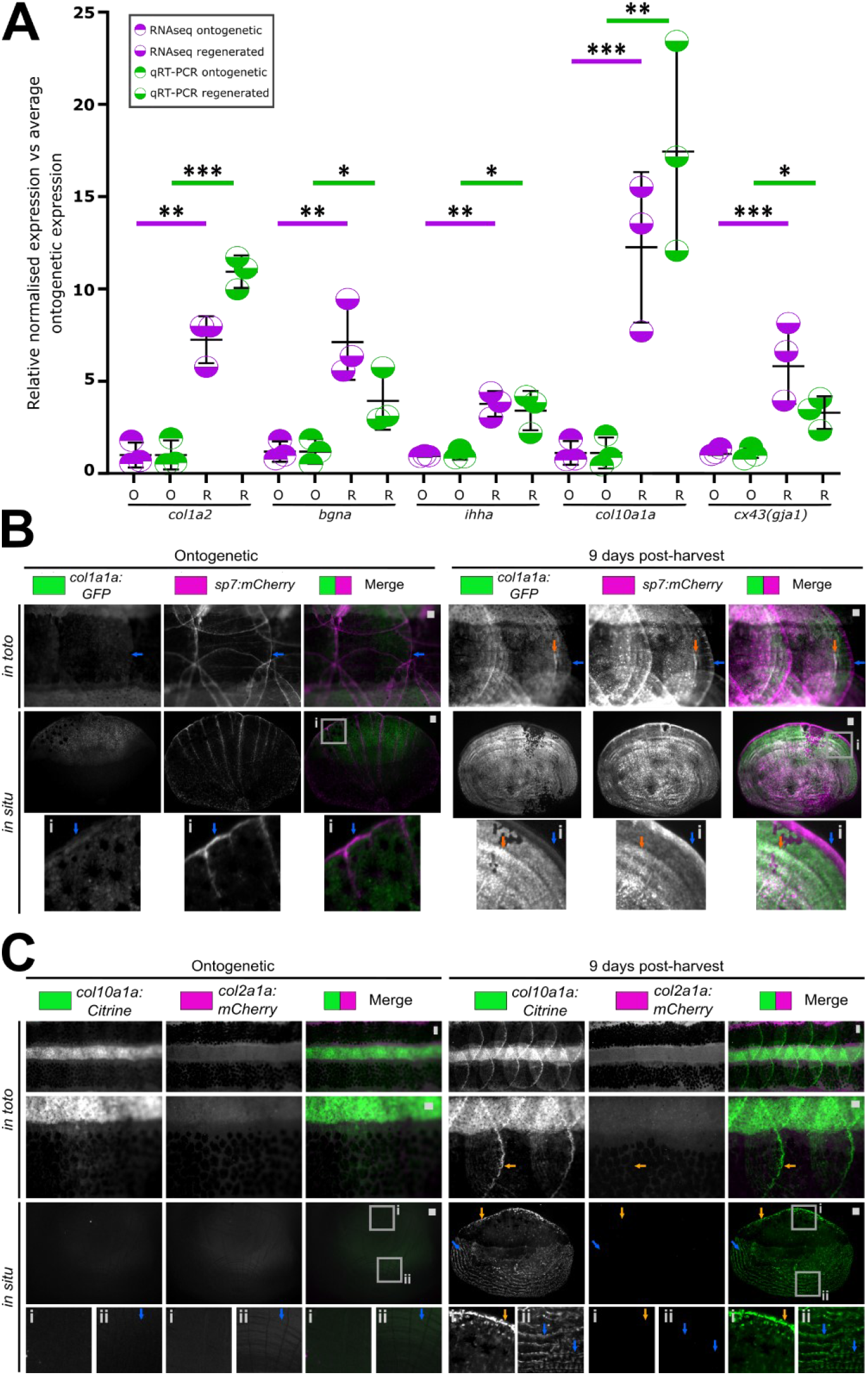
Validation of gene ontology findings. **A)** Relative expression levels of qRT-PCR and transcriptomic analysis of amplicons found in the collagen, proteoglycan and hedgehog signalling networks. **B)** Images of *in toto* and harvested scales (*in situ*) of *col1a1a:GFP* and *sp7:mCherry-NTR* double transgenic fish (n = 4 each condition). Blue arrow indicates the posterior (distal) fringe with high expression of mCherry. Orange arrow points at co-expression of GFP and mCherry. **C)** *In toto* and *in situ* stereomicroscope images of *col10a1a:Citrine* and *col2a1a:mCherry* double transgenic ontogenetic and regenerating scales (n = 4 fish each condition). Orange arrow indicate Citrine signal at the posterior distal edge (inset i) while blue arrow points at a newly forming lateral circulus (inset ii) of the scale. Scale bar: 100 μm.

### Differentially expressed genes are enriched for human orthologues that cause monogenic skeletal disease, and that associate with polygenic musculoskeletal traits and disease

Next, we used a Fisher’s Exact test of independence, in conjunction with the ISDS Nosology and Classification of Skeletal Disorders database (Mortier, Cohn et al., 2019) to show that human orthologues of zebrafish DEGs were 2.8 times more likely to cause monogenic skeletal dysplasia in humans than expected by chance (P=8×10^−11^) (**figure 5A**). Specifically, human orthologues of 47 DEGs resulted in one or more primary bone dysplasia in humans when mutated (**data file S6**). Subgroup analysis revealed further evidence of of enrichment for genes that cause: ‘Osteogenesis Imperfecta and decreased bone density’ (8-fold enrichment, P=7.9×10^−10^), ‘abnormal bone mineralisation’ (7-fold, P=1.6×10^−3^), ‘collagen type 11’ (18-fold, P=3.1×10^−3^), ‘metaphyseal dysplasia’ (6.7-fold, P=7.9×10^−3^), and several others (**figure 5B & data file S6**).

**Figure 5:**
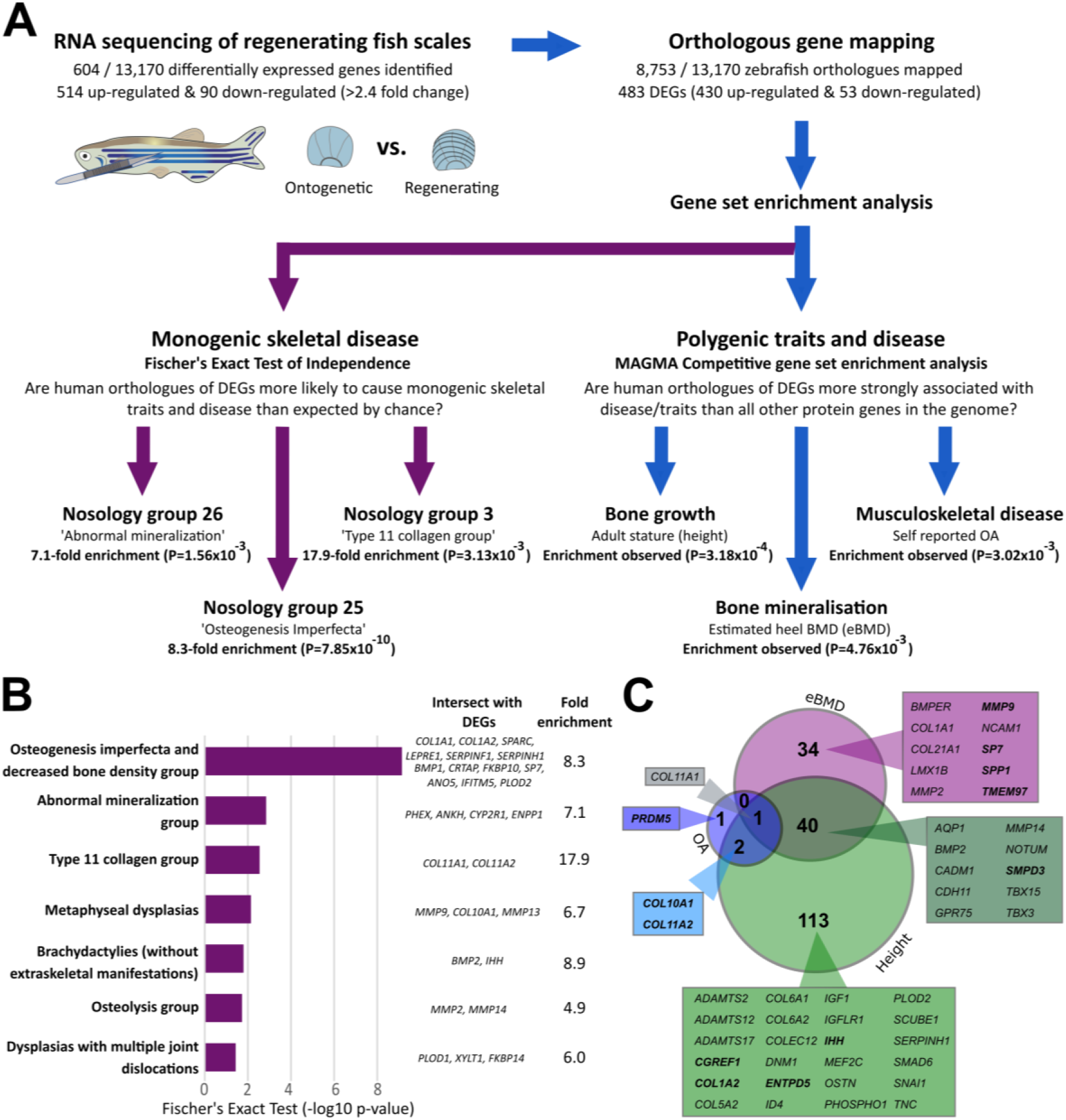
Gene set enrichment analysis strategy showing gene set enrichment of zebrafish DEGs in monogenic skeletal disorders and complex musculoskeletal traits and disease. **A)** Flow diagram summarising gene set enrichment analysis approach showing examples of enriched monogenic nosology groups and polygenic traits using human orthologues of zebrafish DEGs. **B)** Graph showing p-value of monogenic skeletal dysplasia nosology groups found in this study. The fold enrichment and human disease genes and zebrafish DEGs intersect are listed as well. **C)** Venn diagram of gene-based tests of association post-hoc analysis output showing number of associated DEGs with a stringent gene-wide significance threshold of P<2.63×10^−6^. The diagrams show the unique or intersecting genes between estimated bone mineral density (eBMD), osteoarthritis (OA), and height UK-Biobank gene based tests of association. The boxes show selected examples of DEG in each category and bold gene symbols are validated in this study.

MAGMA competitive gene set analysis was used to investigate the relationship between genetic variation surrounding human orthologues of 459 mappable zebrafish DEGs and quantitative ultrasound derived heel bone mineral density (eBMD) measured in 378,484 white European adults (**figure 5A**). Strong evidence of enrichment was observed, suggesting that human orthologues of zebrafish DEGs were on average more strongly associated with the eBMD than all other human protein coding genes in the genome (β=0.20, SE=0.079, P=0.005, **table 2**). In a sensitivity analysis, we adjusted further for the set of orthologues that could not be mapped between fish and humans, and for the set of orthologues that was not expressed in ontogenetic and/or regenerating scales. Adjustment for these potential confounders reduced the magnitude of enrichment by a third, however the set of DEGs remained enriched for BMD associated orthologues (β*=0.14, SE=0.080, P=0.04*). Post hoc permutation analysis suggested that enrichment was not attributable to all DEGs, but rather that a subset of DEGs accounted for most of the enrichment. To further investigate this finding, DEGs were stratified according to whether they were up- or downregulated, and each set was re-analysed. Enrichment for BMD associated orthologues was stronger (β=0.27, SE=0.084, P=0.0006) for the 402 upregulated DEGs and no evidence of enrichment was observed for the 57 downregulated DEGs (β=−0.31, SE=0.23, P=0.91) (**figure S7A**). Subsequent post hoc analysis suggested that enrichment for BMD associated orthologues was attributable to at least ~40% of the upregulated DEGs. We repeated the analysis using height and osteoarthritis (OA) and observed evidence that DEGs were enriched for both outcomes (P<0.005, **figure 5A**, **table 2, figure S7B**). Notably, post hoc analysis revealed that the enrichment for height and OA associated human orthologues was not attributable to all DEGs. Stratified analysis revealed that upregulated DEGs were more strongly enriched for height associated orthologues, whereas the enrichment for OA associated genes remained largely unchanged (**figure S7B, table 2**).

**Table 2:** MAGMA competitive gene set enrichment analysis with p-value of >0.05 in green.

| Model | Trait/Disease | #Genes | BETA | BETA STD | SE | P-value |
| --- | --- | --- | --- | --- | --- | --- |
| **<u>ALL DEGs</u>** |  |  |  |  |  |  |
| Base | eBMD | 459 | 0.20 | 0.03 | 0.079 | 4.76E-03 |
|  | Height | 459 | 0.35 | 0.05 | 0.102 | 3.18E-04 |
|  | Osteoarthritis | 459 | 0.13 | 0.02 | 0.047 | 3.02E-03 |
| Sensitivity | eBMD | 459 | 0.14 | 0.02 | 0.080 | 4.02E-02 |
|  | Height | 459 | 0.24 | 0.04 | 0.103 | 9.96E-03 |
|  | Osteoarthritis | 459 | 0.12 | 0.02 | 0.048 | 6.91E-03 |
| **<u>Upregulated DEGs</u>** |  |  |  |  |  |  |
| Base | eBMD | 402 | 0.27 | 0.04 | 0.084 | 5.61E-04 |
|  | Height | 402 | 0.39 | 0.06 | 0.109 | 1.45E-04 |
|  | Osteoarthritis | 402 | 0.14 | 0.02 | 0.050 | 3.26E-03 |
| Sensitivity | eBMD | 402 | 0.21 | 0.03 | 0.085 | 6.57E-03 |
|  | Height | 402 | 0.29 | 0.04 | 0.110 | 4.56E-03 |
|  | Osteoarthritis | 402 | 0.12 | 0.02 | 0.051 | 7.05E-03 |
| **<u>Downregulated DEGs</u>** |  |  |  |  |  |  |
| Base | eBMD | 57 | -0.31 | -0.02 | 0.227 | 9.13E-01 |
|  | Height | 57 | 0.01 | 0.00 | 0.292 | 4.86E-01 |
|  | Osteoarthritis | 57 | 0.07 | 0.00 | 0.131 | 2.89E-01 |
| Sensitivity | eBMD | 57 | -0.38 | -0.02 | 0.228 | 9.53E-01 |
|  | Height | 57 | -0.10 | -0.01 | 0.292 | 6.38E-01 |
|  | Osteoarthritis | 57 | 0.06 | 0.00 | 0.131 | 3.29E-01 |
| Key: described in supplemental data file S7 |  |  |  |  |  |  |

Finally, one advantage of MAGMA is that it first performs gene-based tests of associations between each human protein coding gene in the genome, and the outcome of interest (see supplementary methods for more detail). Consequently, once enrichment is observed we can prioritise orthologues of DEGs for functional validation by comparing their relative strength of association with BMD, height and/or OA (**figure 5C** and **data file S7**). Using this approach, we identified human orthologues of *spp1* and *col11a2* that were both differentially upregulated in regenerating fish scales, and showed different patterns of association with all three traits. Specifically *SPP1* was robustly associated with BMD (P=4×10^−15^), but not height (P=0.16) or OA (P=0.16), whereas *COL11A2* was robustly associated with height (P=4.1×10^−13^) and OA (P=3.6×10^−7^), but did not meet the stringent gene-wide significance threshold of P<2.63×10^−6^ for BMD (i.e. P=2.5×10^−3^). Given these contrasting patterns of association, together with the availability of the corresponding mutant zebrafish lines, we chose to further investigate *spp1* and *col11a2* expression and establish whether mutant fish developed skeletal abnormalities that were consistent with the above mentioned human genetic associations.

### Zebrafish mutants of differentially expressed genes show skeletal phenotypes consistent with their expression profile and with human genetic associations

Human orthologues of DEGS that had unique, or pleiotropic associations with eBMD, height and/or OA were identified (**figure 5C**), and qRT-PCR was used to confirm that the corresponding DEGs (including *spp1* and *col11a2*) were differentially expressed (**figure 6A**). We further assessed transgenic reporter expression of *spp1* (*spp1:mCherry*) to determine gene expression localisation. The expression pattern of *spp1:mCherry* was exclusively localised at the distal rim of the ontogenetic scale, similar to *sp7:GFP* positive osteoblast localisation (**figure 6B**). Interestingly, these *sp7* positive (sub)marginal cells at the rim are classed as more mesenchymal osteoblasts involved in *de novo* bone formation in ontogenetic scales (Iwasaki, Kuroda et al., 2018). In regenerating scales, *sp7* expression was seen more broadly but elevated at the posterior edge which was followed proximally by an increased *spp1* signal with reduced *sp7* expression in the same area (**figure 6B**). The width of *spp1* expression was increased compared to ontogenetic scales, implying an expansion of its activity during bone growth and mineralisation in the regenerating scale (**figure 6C** and **6D**).

**Figure 6:**
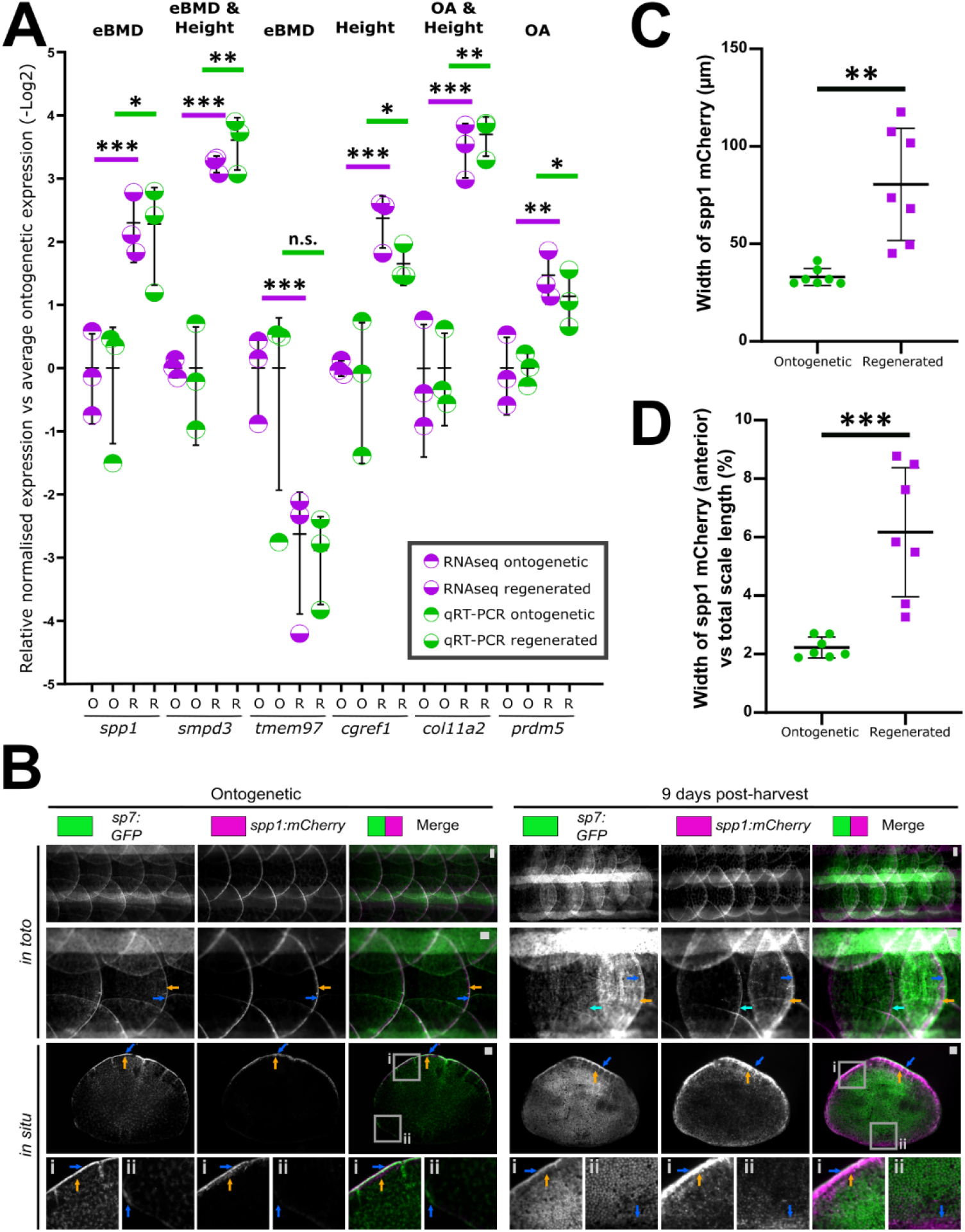
Validation of genes associated with polygenic traits in humans. **A)** Q-RTPCR and RNA-seq expression levels (relative to average ontogenetic expression) of selected DEG found in each category. All but one (*tmem97*) reached statistical threshold of p<0.05. **B)** Images of scales from *in toto* and harvested scales of *sp7:GFP* and *spp1:mCherry* double transgenic fish (ontogenetic: n = 5, regenerating: n = 4 fish). For overview and inset i, blue arrows indicate high GFP signal and orange the mCherry signal at the posterior edge (inset i) while light blue arrow indicates an ontogenetic scale next to a regenerating scale. Inset ii shows anterior region of the scale with opposite GFP and mCherry signals (blue arrow) in ontogenetic and regenerating conditions. **C)** Quantification of width of the mCherry signal at the posterior edge from harvested scales (between blue and orange arrows in panel C). **D)** Width of mCherry signal normalised by scale length (anterior to posterior). Scale bar: 100 μm.

We next measured axial skeleton length from 3D micro-CT renders of *col11a2*^*Y228X/Y228X*^ and *spp1*^*P160X/P160X*^ mutant zebrafish. This showed significant reduced length in *col11a2* mutants whilst *spp1* mutants exhibited a milder tendency (**figure 7A**). Additionally, we segmented the lower jaw (mandibular arch) and caudal vertebrae of these 3D micro-CT renders and measured several histomorphological parameters using element landmarks as set out in **figure S8A-B**. These measurements of the lower jaw demonstrated that *col11a2* fish have reduced width and length in the lower jaw whereas *spp1* fish did not show significant changes (**figure 7B**). When we calculated the ratio between length and width, both mutant lines showed a mild alteration in lower jaw element proportion (**figure 7C**). Interestingly, in contrast to wildtype and *spp1* mutants, the *col11a2* mutants exhibited altered joint shape consistent with the predicted OA phenotype in adulthood and as seen in larval jaws previously (**figure 7D**)(Lawrence, Kague et al., 2018). Measurements of the centrum, neural arch, and haemal arch of the vertebra did not show significant changes in either mutant line (**figure 7E-G, figure S8C-G**) These histomorphological data show that indeed *col11a2* fish showed bone growth defects while *spp1* mutants did not.

**Figure 7:**
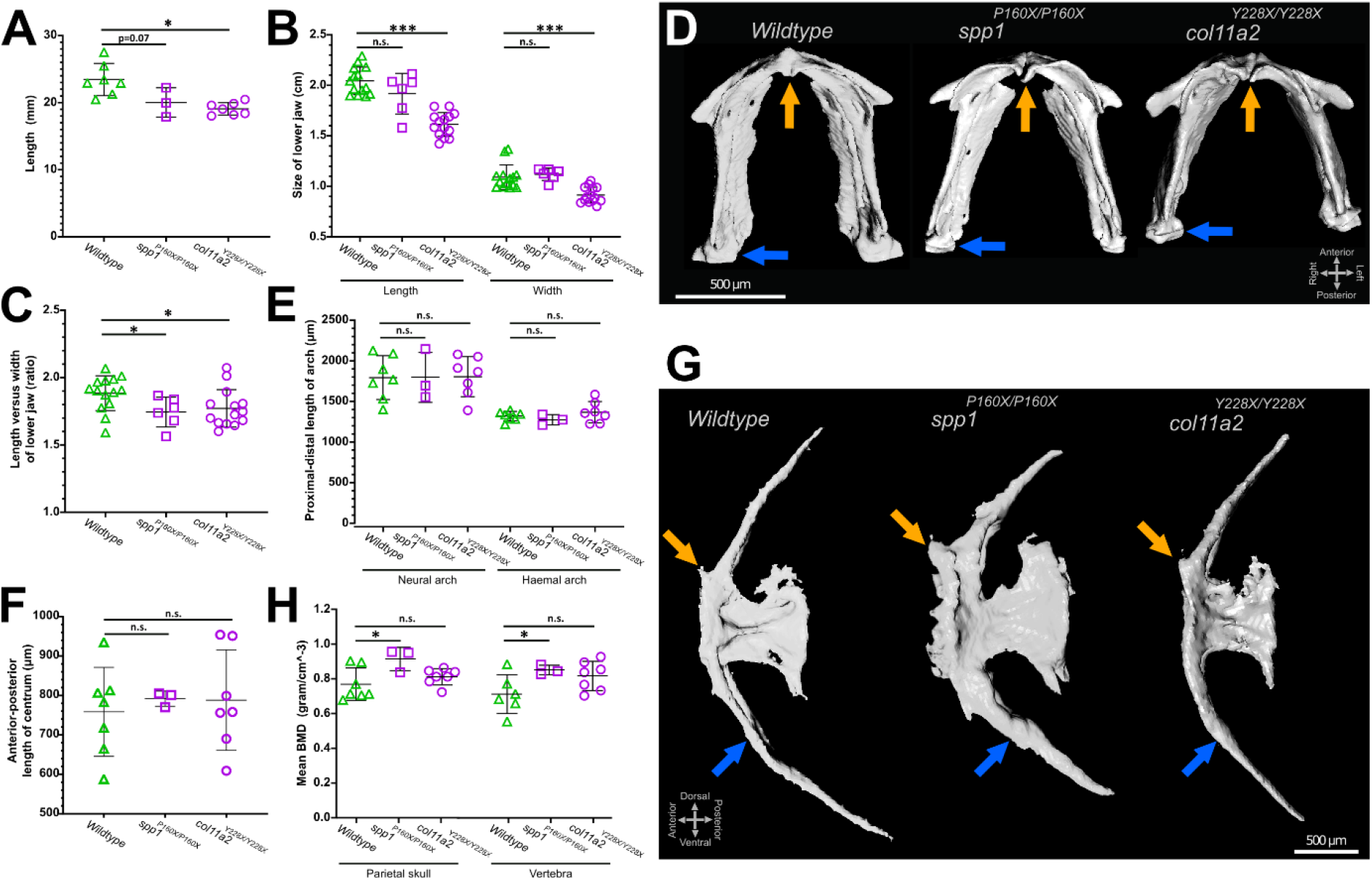
Histomorphology and bone mineral density measurements on 3D micro-CT images of *col11a2*^*Y228X*^ and *spp1*^*P160X*^ homozygous fish revealed altered bone structures. **A)** Axial skeleton length was reduced in *col11a2* mutants whereas *spp1* mutants showed a mild reduced tendency. **B)** Quantification of lower jaw size. **B)** Calculation of lower jaw element proportions. **D)** Ventral view of the segmented lower jaw images. Orange arrow indicates anterior mandibular arch joint and blue arrow shows mandibular arch – palatoquadrate (not visible) joint. **E)** Quantification of the vertebral arches’ length. **F)** Total length of the anterior facet of the vertebral centrum. **G)** Lateral view of segmented images of the first caudal vertebra showing the anterior facet (orange arrow) and haemal arch (blue arrow). **H)** Mean BMD calculations of dermal and notochord sheath derived bone.

To test the prediction that *spp1* mutants would have abnormal BMD, we quantified mean volumetric BMD from sites formed via different modes of ossification: the parietal plate of the cranium and the first caudal vertebra as these are formed by intramembranous ossification and by ossification directed by the notochord sheath, respectively. Consistent with our prediction, we observed an increase in BMD for *spp1* mutants in both elements whereas no evidence of altered BMD was observed for *col11a2* mutants (**figure 7H**). Interestingly, we observed a physical thickening of the vertebral arches and anterior shaft of the vertebral centrum in *spp1* mutants consistent with the relationship seen with the increased BMD (**figure 7G** and **figure S8E**). Taken together, our data suggest that genes that are differentially expressed during scale regeneration play a role in wider regulation of skeletal homeostasis at other skeletal sites in the zebrafish, and that mutants in these genes have phenotypes that are largely consistent with those observed in humans.

## Discussion

In this study we define, for the first time, the transcriptome of ontogenetic and regenerating zebrafish scales and show that DEGs are enriched for biological pathways involved in osteoblast mediated bone formation, many of which are conserved in humans. By integrating this information with large-scale human genetic association studies, we show that human orthologues of DEGs were likely to play a role in the pathogenesis of monogenic and polygenic human MSK disease. Our findings suggest that zebrafish scale regeneration has the potential to help us better understand biological function and pathways relevant to human skeletal health and disease.

Transcriptomic profiling revealed that regenerating scales predominantly upregulate gene expression of many osteoblast genes, correlating with the high abundance of metabolically active osteoblasts. The high expression of genes involved in ECM deposition, principally genes regulating collagen synthesis, processing, and deposition fits with the regeneration of the collagen-rich matrix of the bony scale. We demonstrated that while *col1a1a* is expressed throughout the regenerating scale it is largely excluded from the leading edge labelled by *sp7.* By contrast, *col10a1a* is strongly expressed in leading edge concomitant and newly forming lateral circuli that are associated with thickening of the calcified matrix in accordance with the role of type X collagen as an early marker of ossifying tissues in the zebrafish (Debiais-Thibaud, Simion et al., 2019, Eames, Amores et al., 2012, Hammond & Schulte-Merker, 2009). Amongst DEG collagens, *col11a2* was one of the highest DEGs in this profile. Interestingly, type XI collagen is more frequently associated with cartilage matrix formation; interacting with type II collagen to regulate spacing and nucleation of fibrils in the ECM (Gregory, Oxford et al., 2000, Li, Lacerda et al., 1995). As well as the collagens themselves, matrix processing genes coding for MMPs were also upregulated. MMPs are important to breakdown the dense type I collagen matrix and this process is crucial for tissue growth, shape, release and distribution of signalling molecules, and cell migration (Page-McCaw, Ewald et al., 2007). We have previously shown that *mmp9* is upregulated during scale regeneration and *mmp9* expression was observed adjacent to newly deposited matrix and TRAP positive cells (de Vrieze et al., 2011).

Moreover, we observed transcriptional upregulation of ECM proteoglycans that bind collagens, such as biglycan *(bgna/b*), which are important in regulating collagen nucleation and remodelling (Douglas, Heinemann et al., 2006). Interestingly, biglycan has been shown to modulate ECM accessibility and regulate diffusion of signalling molecules, such as Bmp4, regulating bone strength in mice (Chen, Fisher et al., 2004, Xu, Bianco et al., 1998). Related to the abundant presence of collagen related factors, cell adhesion genes, in particular integrin signalling, were enriched and connected to the collagen and MMP networks. Cell adhesion and the ECM are interlinked and both must be tightly regulated during rapid growth (Zeltz & Gullberg, 2016), such as osteogenic regeneration of the scale. Cell adhesions are heavily modulated to allow cell rearrangements and tissue mechanics in regenerative tissue expansion contexts. For example, in a mammalian epidermal wound healing response it is crucial for epithelial regeneration (Mosaffa, Tetley et al., 2020).

Pathway analysis revealed signalling pathways related to ECM formation and expansion, such as integrin, HH and IGF. The absence of enrichment (note these were also not under-represented) of Wnt, Bmp, and Fgf pathway genes is likely to be associated with the timing of the transcriptomic profile (9 dph). For example, it has been shown that Wnt and Fgf signalling are required for the initiation of scale regeneration, with inhibiting Fgf or canonical Wnt signalling leading to a failure to form scales (Aman, Fulbright et al., 2018). HH signalling controls morphogenesis of the scale over a prolonged time period, where the HH ligand secreted from the epidermis regulates the rate of ECM deposition by scale osteoblasts (Aman et al., 2018, Iwasaki et al., 2018).

Gene ontology analysis suggested that transcriptomic profiles of regenerating zebrafish scales were enriched for biological pathways involved in bone formation and mineralisation. These pathways are largely conserved in humans, prompting us to investigate whether DEGs were enriched for human orthologues that cause human monogenic skeletal disorders, and that associate with polygenic MSK traits and disease. Gene set analysis involving human genetic association studies complemented findings from our gene ontology analysis, suggesting that DEGs were most strongly enriched for human orthologues that cause monogenic disorders that were broadly characterised by abnormal or decreased bone mineralisation, and separately, abnormal cartilage formation and bone growth. Further analysis involving human polygenetic disease traits provided additional evidence that upregulated (but not downregulated) DEGs, were enriched for human orthologues associated with bone mineralisation as captured by heel bone ultrasound. Importantly, post hoc permutation analysis revealed that although the set of upregulated DEGs was strongly enriched, it was likely that enrichment was not attributable to all DEGS, but rather a nested subset (or subsets) of genes that corresponded to one or more biological pathway(s) that regulates bone homeostasis in humans. A possible explanation is that DEGs were identified using bulk RNA-sequencing, a method that captures a heterogenous mixture of transcriptome signature profiles that define different cell types, states, and their biological pathways, some of which may or may not be involved in bone mineralisation during scale regeneration. Importantly, the small sample size of our transcriptomic study also limited our ability to identify all DEGs and their associated biological pathways. However, as the scale contains a heterogeneous cell population, this is the most likely cause of not identifying the nested subsets of DEGs responsible for the enrichment. Hence, future studies may benefit by focusing on transcriptomic profiling of separate cell populations from scales at different stages of regeneration to determine their expression profiles to dissect the different pathways responsible for osteo-anabolic growth. This could be achieved by, for example, using fluorescent automated cell sorting (FACS) to isolate various populations using cell specific transgenic reporters (e.g. *sp7:GFP*) or even single cell RNA-seq technologies to elucidate the transcriptional state of all different cell type populations in the regenerating scale.

Given that DEGs appeared enriched for human orthologues associated with human MSK traits and disease, we chose to perform detailed skeletal phenotyping on *spp1* and c*ol11a2* zebrafish mutants to investigate whether they developed skeletal abnormalities that were consistent with their gene-based associations with height, eBMD and/or OA. Both genes were upregulated DEGs in regenerating scales and human orthologues of these genes showed contrasting patterns of association with human disease traits (i.e. *SPP1* robustly associated with eBMD only, and *COL11A2* robustly associated with height and OA, and suggestively associated with *BMD).* Mutant fish developed skeletal abnormalities largely consistent with those predicted by the human genetic association studies. Notably, in humans, mutations in *COL11A2* lead to Stickler syndrome, a condition associated with craniofacial dysplasias, and joint abnormalities that lead to premature OA (Couchouron & Masson, 2011). No clear human phenotype for *SPP1* (osteopontin) has been defined yet, but it plays a complex role in regulating repair and regeneration of multiple tissues in vertebrates. We show that *spp1:mCherry* was up-regulated in the regenerating scale adjacent to pre-osteoblastic sp7+ cells which is consistent with previous reports that expression of *spp1* is up-regulated during bone remodelling in both zebrafish fins and mammals (Morinobu, Ishijima et al., 2003, Sousa, Valerio et al., 2012, Terai, Takano-Yamamoto et al., 1999). Moreover, *spp1* mutants have elevated BMD in the cranial and axial endoskeleton, in line with murine studies where loss-of *Spp1* leads to enhanced mineral content in some parts of trabecular bone (Boskey, Spevak et al., 2002). Mouse studies have shown that *Spp1* determines mineralisation rate in the skeleton through regulating *Phospho1* expression (Holm, Gleberzon et al., 2014, Yadav, Huesa et al., 2014). For example, Osteopontin expression is detected during cranial suture closure in both mice and zebrafish, suggesting a role in appropriate timing of pre-osteoblast differentiation (Kim, Lee et al., 2003, Morinobu et al., 2003, Topczewska, Shoela et al., 2016). Outside of its function in skeletal tissue, mammalian *Spp1* is functionally diverse, as it promotes angiogenesis and its expression is also triggered during cutaneous wound healing to control the rate of repair (Dai, Peng et al., 2009, Mori, Shaw et al., 2008). As *Spp1* is a component of the ECM and possesses integrin binding domains, it can bind various integrins to modulate cell adhesion to the collagen matrix important during i.e. tissue growth (Lamort, Giopanou et al., 2019). As we see high *spp1:mCherry* expression adjacent to *sp7*+ pre-osteoblasts in the regenerating scale, it implies that *spp1* could play a role in differentiation of these pre-osteoblasts and therefore timing of scale mineralisation and the association with human eBMD is suggestive of a conserved function in control of BMD across species.

Since the prevalence of diseases with pronounced bone fragility phenotypes is increasing due to an ageing population there is a need to discover and rapidly test new bone growth candidate genes that could act as drug targets. These multi-factorial diseases have complex genetic and physiological underpinnings that are not well understood, and as human bone samples are hard to obtain there is a strong demand for models to study bone pathophysiology. The regenerating zebrafish scale could therefore function as an additional model to study and discover osteo-anabolic factors relevant to human skeletal diseases. The relative abundance of teleost scales (around 200 per fish) and their amenability for imaging make them an ideal model for regenerative studies, and unlike caudal fins, scales can be treated as independent bone units, each harbouring their native complex tissue environment. Scales can be cultured *ex vivo* in a semi-high throughput multi-well format, offering an avenue to test compounds that could complement established tissue culture and *in vivo* pharmacology studies (Bergen et al., 2019, de Vrieze et al., 2015). By using a multi-disciplinary approach we: i) described a transcriptomic profile of regenerating zebrafish scales, ii) showed that DEGs are enriched for biological pathways involved in bone formation, and for genes associated with MSK disease in human populations and, iii) studied mutant zebrafish for two DEGs and observed skeletal abnormalities in both fish that were consistent with results from human genetic association studies. In conclusion, we show that integrative analysis involving zebrafish transcriptomics and human genetic association studies is feasable, and that future studies involving zebrafish scales have potential to better our understanding of bone formation in general, and how defects in this complex process contribute to musculoskeletal disease pathogenesis in humans.

## Material and Methods

### Zebrafish husbandry, mutant and transgenic lines

Zebrafish were maintained under normal husbandry conditions(Alestrom, D'Angelo et al., 2020). Experiments were locally ethically reviewed (by AWERB at University of Bristol (Bristol, UK) (UoB) and Radboud University (Nijmegen, NL) (RU) respectively) and performed under UK Home Office project licence 30/3408 (UoB) or RU-DEC2014-059 (RU).

Wildtype AB/TL (UoB) and AB (RU) strains were used. Mutant lines (AB/TL background, UoB) have been previously described; *col11a2*^*sa18324*^ carries a nonsense mutation causing a premature stop codon at tyrosine position 228 (ENSDART00000151138.3), henceforth called *col11a2*^*Y228X*^ (Lawrence et al., 2018) and *spp1*^*CGAT327-330del*^ carrying a deletion leading to a frameshift resulting in a premature stop codon nonsense mutation at proline position 160 (ENSDART00000101261.6), henceforth called *spp1*^*P160X*^ (Bevan, Lim et al., 2020). Transgenic lines are listed in the **Supplemental Methods**.

### Elasmoid scale harvesting and imaging

Anaesthetised fish (0.05% (v/v) tricaine methanesulfonate (MS-222) (UoB), 0.1% (v/v) 2-phenoxyethanol (RU)) were put on a wet tissue containing system water and anaesthetic, and scales plucked under a microscope with a watchmaker’s tweezers from the midline of the lateral flanks near the dorsal fin. Fluorescent microscope images of flanks (*in toto*) and single harvested scales (*in situ*) were acquired on a fluorescent stereomicroscope (Leica Microsystems, Germany), using 2x, 4x, and 8x magnification.

Adult fish were immersed in 40 μM Calcein (Sigma-Aldrich, cat# 154071-48-4) Danieau’s buffer solution (pH 7.4) for two hours and washed in system water for at least 15 minutes prior to imaging.

#### Alkaline phosphatase staining

Scales were collected in ALP buffer (100 mM Tris-HCl (pH 9.5), 100 mM NaCl, and 50 mM MgCl_2_) and stained in ALP buffer containing 2% (v/v) NBT/BCIP (Sigma, cat# 11681451001). After a brief wash in deionised water, scales were mounted on a microscope slide containing 10 % (w/v) Mowiol^®^ 4-88 (Sigma-Aldrich cat# 9002-89-5) in 25% (v/v) glycerol solution. Images were taken on an upright microscope (Leica).

### RNA isolation, RNA sequencing, transcriptomic mapping and analysis

Approximately 40 scales were collected from a standardized area on the left flank of 1-year old male zebrafish; the area that extends from just behind the operculum to the implant of the dorsal fin. The area may included multiple rows of scales of similar size and shape. Total RNA was isolated from ontogenetic and regenerating scales (n=3 fish per group, RU) by using Trizol (Invitrogen) for RNA-sequencing (RNA-seq) and downstream qRT-PCR testing.

RNA-seq was performed by ZF-GENOMICS (Leiden, NL) and involved quality control of total RNA libraries on 2100expert bioanalyzer (Agilent) that resulted in RIN scores of >9.2. Illumina RNAseq library preparation involved standard 6 nucleotide adaptor ligation. Paired single read 1 x 50 nucleotide runs (10 million reads; 0.5 Gb per sample) were performed on an Illumina Hiseq2500 system.

#### Transcriptome Mapping and Differential Expression Analysis

Raw reads were mapped to the GRCz11 primary assembly (Ensembl version 99) using STAR (version STAR_2.5.4b) software pipeline (Dobin, Davis et al., 2013). The read count table for all genes mapped was obtained from the mapping step and filtered to leave out lowly expressed genes by only keeping genes that had at least 5 mapped reads over all samples. The differential gene expression analysis was performed using the R-package DESeq2 (version 1.28.1)(Love, Huber et al., 2014) including the medians of ratio normalisation step to account for the bias in sequencing depth/coverage and RNA composition of samples. For determining DEGs, we used a threshold of 1.25 log2 fold (2.4-fold) change and a false discovery rate (FDR) of <0.05. Further details are in the **Supplemental Methods**.

For downstream analyses, genes were classed as ‘expressed’ in scales when all three ontogenetic and regenerating scale samples produced a >5 normalised read count (background gene list). Arbitrary threshold for differential expression was set at ±1.25 log_2_ fold and an adjusted p-value (padj) of ≤0.05 (DEG list).

### Gene ontology enrichment and STRING network analysis

DEG and background expression gene lists’ gene symbols were uploaded to GOrilla (Eden, Navon et al., 2009) using *Danio rerio* and ‘two unranked lists of genes’ as settings. The hierarchical images and Microsoft Excel files were used for figure making. Ensembl IDs of DEGs were analysed with an ‘Overrepresentation Test’ (Released 20200407) using Fisher’s exact test and False Discovery Rate correction in PANTHER Gene Ontology (release 2020-06-01), and PANTHER ‘Pathways’ and ‘Reactome pathways’ (PANTHER version 15.0) software (Mi, Muruganujan et al., 2019).

For STRING (v11) gene network analysis (Szklarczyk, Gable et al., 2019), the DEG set (zebrafish gene symbols) was uploaded and interaction score (high, ≥0.7 or medium ≥0.4), number of interactions (one shell with max. 10 interactions), and active interaction sources (all were on) were set. More details are in the **Supplemental Methods**.

### Gene set enrichment analysis involving monogenic and polygenic skeletal traits and disease

In conjunction with ISDS Nosology and Classification of Skeletal Disorders database (Mortier et al., 2019), evidence of enrichment for human monogenic skeletal disease causing genes either total or within each individual nosology-defined skeletal disorder groups) was performed using Fisher's Exact Test of Independence as described in detail in the **Supplemental Methods** and (Youlten, Kemp et al., 2020). Enrichment for polygenetic traits and disease was investigated using MAGMA competitive gene set analysis. In the first stage zebrafish – human orthologues of all protein coding genes were mapped. In the second stage, the strength association between each human protein coding gene and eBMD and height and any form of self-reported or hospital defined OA was evaluated separately using the weighted average of 3 different gene-based tests of association. Gene based tests of association were performed using high quality genome-wide imputed genetic data (~12 million SNPs, INFO > 0.9, MAF > 0.05%) from 378,484 white European adults from the UK-Biobank Study with eBMD and height measures. For the analysis involving OA, publically available genome-wide association summary results statistics were used instead of genome-wide imputed genotyping data (Tachmazidou, Hatzikotoulas et al., 2019). In the final stage, competitive gene set analysis was used to compare the mean strength of association of human orthologues of zebrafish DEGs with each trait / disease, to the mean strength of association all other genes, and evidence of enrichment was quantified using a one-sided test of statistical significance.

Sensitivity analysis was performed correcting for the set of human genes that could not be mapped between human and zebrafish, and the set of human orthologues that was not expressed in zebrafish. Finally, post hoc permutation analysis was performed for analyses that were suggestive of enrichment. A detailed description of the methods is presented in the **Supplemental Methods**.

### Quantitative real-time PCR

500ng of total RNA was treated with DNase (1 unit) and reverse-transcribed (random hexamer primers) with SuperScript II (Invitrogen 100 units). iQ SYBR Green Supermix (Biorad) containing 350 nM primer and cDNA was used for amplification (primers and PCR conditions in **Supplemental Methods**). Relative expression was calculated based on a normalisation index of two reference genes: *eef1a1l1* and *rpl13*. For comparison of qRT-PCR and RNA-seq expression of amplicons, the average of ontogenetic normalised expression (qRT-PCR) or read count (RNA-seq) were taken and every individual value was compared to the average ontogenetic expression (e.g. read count regenerating scales individual 2 / average read count ontogenetic). The p-values presented in the figures were derived from a two-tailed t-test (qRT-PCR) and p-adjusted (padj) from the DESEQ2 analysis (RNA-seq).

### Micro-Computed Tomography and BMD calculations

MicroCT was performed as previously described (Kague et al., 2019), with BMD calculated as previously described (Stevenson et al., 2017). Briefly, adult WT sibling, *col11a2*^*Y228X*^ and *spp1*^*P160X*^ mutant zebrafish (1 year old) were fixed, dehydrated to 70% EtOH and scanned at 21 μm voxel size (scan settings 130 kV, 150 μA, 0.5-s exposure, 3141 projections). Images were reconstructed using NRecon software (Version 1.7.1.0), with dynamic ranges calibrated against a scan of hydroxyapatite phantoms (0.25 g.cm^−3^ and 0.75 g.cm^−3^) scanned with identical parameters. For BMD calculations Aviso software (Avison2020.2; Thermo Fisher Scientific) was used to isolate pixel greyscale values for the skull (parietal) and vertebrae (vertebrae 11-13) and calibrated against the greyscale values of known hydroxyapatite density phantoms (0.25 g.cm^−3^ and 0.75 g.cm^−3^) to estimate mean BMD values and their standard deviations. Histomorphological assessment of segmented jaw and vertebrae (11-13) elements were determined by measuring the width, length, and depth of the elements (N.B. left and right side of jaw elements were considered as independent measurements) in AVIZO between element landmarks as shown in **figure S7A** and **7B**.

### Statistical testing

All graphs and data of downstream RNA-seq functional data were analysed in GraphPad Prism (v 8.4.3) and were subjected to variance comparisons (F test) to determine distribution of the data prior to performing an unpaired t-test comparing ontogenetic against regenerating or wildtype against mutant conditions. P-values in figures as follows: n.s. >0.05, * ≤0.05, ** ≤0.01, *** ≤0.001. RNA-seq, GO, STRING pathway, and GSEA statistical analyses are described in their relevant sections.

## Acknowledgements

We would like to thank our funding sources: D.B. and C.H. received Fellowship funding from Versus Arthritis (22044 and 21937 respectively). J.M. and G.F. were supported by the subsidy programme Smartmix (SSM06010) of the Dutch Ministries of Economic Affairs and Education, Culture and, Science. J.P.K was funded by a University of Queensland (UQ) Development Fellowship (UQFEL1718945), a National Health and Medical Research Council (Australia) Investigator grant (GNT1177938) and project grant (GNT1158758). M.L. is supported by a UQ Research Training Scholarship and the Commonwealth Scientific and Industrial Research Organisation Postgraduate Top-Up Scholarship. R.R. and R.J.R. were supported by the BHF Oxbridge Centre of Regenerative Medicine (RM/17/2/33380) and a BHF Intermediate Fellowship to R.J.R. (FS/15/2/31225). This research has been conducted using the UK Biobank Resource (accession ID: 53641). Lastly, we would like to thank Mathew Green and Pablo Peon Garcia for fish care at UoB, and Tom Spanings and Antoon van der Horst for fish care at RU.

## Author contributions

D.B., and J.M. conceptualised and initiated the project. D.B., Q.T., J.P.K., CH., and J.M. designed the experiments. D.B., Q.T., A.S, J.Z., E.N., J.P.K., M.L., S.Y., C.H., and J.M. performed the experiments and analysed data. E.Z, S.Y., J.P.K, R.R. and R.J.R. provided reagents or access to datasets. D.B., J.P.K., C.H., and J.M. supervised the project. D.B., G.F. J.P.K., CH., and J.M. acquired funding for the project. D.B. prepared manuscript figures. J.P.K, E.Z., S.Y, P.I.C., R.R., G.F. and R.J.R. provided expertise. D.B., J.P.K., C.H., and J.M. co-wrote the first version of the manuscript. All authors revised the first version and approved the contents of the final version of the manuscript.

## Conflict of interest and ethics statement

All authors declare no conflict of interest. Human genetic and animal research that is presented in this work has been conducted with ethical approval as described in Material and Methods section.

## Data availability

RNA-sequencing FASTQ files are available in ENA (https://www.ebi.ac.uk/ena) under accession number PRJEB39971.

