## Supplementary material for "The transcriptome of regenerating zebrafish scales identifies genes involved in human bone disease": Supplemental Methods.docx (from another version of this preprint: version 1, 2020-10-08)

### Supplemental Material and Methods, Bergen et al.

#### Zebrafish transgenic lines

Transgenic lines were kept in an AB/TL background and full nomenclature as follows: *spp1:mCherry [TgBAC(spp1:mCherry)^SN:374^* (Bevan, Lim et al., 2020)], *col2a1a:mCherry* [*Tg(Col2a1aBAC:mCherry)* (Hammond & Schulte-Merker, 2009)], *sp7:GFP* [*Tg(Ola.sp7:NLS-GFP)* (DeLaurier, Eames et al., 2010)], *sp7:mCherry* [*Tg(osterix:mCherry-NTRo)^pd46^* (Singh, Holdway et al., 2012)], *col1a1a:GFP* [*Tg(col1a1a:EGFP)^zf195tg^* (Kague, Gallagher et al., 2012)], and *col10a1a:Citrine* [*TgBAC(col10a1a:Citrine)* (Mitchell, Huitema et al., 2013)].

#### Transcriptome Mapping

The genomic FASTA sequence, cDNA and GTF annotation files of the latest release (version 99) of reference genome assembly of zebrafish were downloaded from Ensembl (Genome Reference Consortium Zebrafish Build 11). Quality control of the raw reads and adapter trimming was performed by the sequencing vendor. The reference genome was indexed using the genomeGenerate function of STAR (version STAR_2.5.4b) (Dobin, Davis et al., 2013) with the following command and parameters:

*STAR	--runMode genomeGenerate	--runThreadN 16	--genomeDir Genome_Ref	--genomeFastaFiles GCF_000002035.6_GRCz11_genomic.fna	--sjdbGTFfile GCF_000002035.6_GRCz11_genomic.gtf	--sjdbOverhang 50*

Next, mapping of the paired reads (50 bps) from samples to the reference genome was performed using shell scripting and STAR aligner with the following command and parameters:

*for i in {1..2}; do r1="Read1.fastq" r2="Read2.fastq" STAR --genomeDir* *Genome_Ref --runThreadN 328 --readFilesIn $r1 $r2 --outFileNamePrefix Sample_ONT$i.aligned --outSAMtype BAM SortedByCoordinate --quantMode Sample_ONT1_TranscriptomeSAM; done*

The raw read counts were directly obtained from the –quantMode functionality of STAR and genes with zero counts over all samples were removed using R programming language (R version 3.6.3) We obtained a uniform read mapping across all of our samples, with more than 70% of uniquely reads mapped, therefore no further filtering of the data was required.

#### Differential Expression Analysis

The read count table for all gene mapped was obtained from the mapping step and filtered to leave out lowly expressed genes by only keeping genes that had at least 5 mapped reads over all samples. The differential gene expression analysis was performed using the R-package DESeq2 (version 1.28.1) (Love, Huber et al., 2014) including the medians of ratio normalization step to account for the bias in sequencing depth/coverage and RNA composition of samples. Using the built-in function *DESeqDataSetFromMatrix*, the counts data was loaded with the following parameters:

*fitType=local, test=Wald*, and the batch effect correction included in the design formula.

#### Linear Variance Analysis

The variance of gene expression across all samples (ontogenetic and regenerative scales) was calculated using the R-package matrixStats (version 0.56.09) and an in-house R script (R version 3.6.3) (see SUPPLEMENTARY R SCRIPT below).

#### Principal Component Analysis

The clustering of the samples based on their variance in gene expression was performed using the R-package pcaGoPromoter (version 3.11) and an in-house R script (R version 3.6.3) (see SUPPLEMENTARY R SCRIPT below).

#### Volcano plot

Data from the background expression list (**supplemental data file 1**) in Excel was converted to a tab delimited text file containing -log_10_ transformed ‘padj’ values (’minus.log10.padj‘) and a volcano plot was plotted in R (see SUPPLEMENTARY R SCRIPT below).

#### R scripts for Volcano plotting, Linear Variance and Principal Component Analyses

***##Linear Variance Analysis R functions***

```
library(matrixStats)
variance_regression <- function(as.matrix(Normalized-expression-matrix-from-STAR)){
  dis <- data.frame()
  for (i in 1: nrow(as.matrix(Normalized-expression-matrix-from-STAR)){
    dis[i,1] <- rowMeans(as.matrix(Normalized-expression-matrix-from-STAR)[i, , drop=FALSE])
    dis[i,2] <- matrixStats::rowVars(as.matrix(Normalized-expression-matrix-from-STAR)[i, , drop=FALSE])
  }

  plot(log2(dis[,1]+1), log2(dis[,2]+1),pch=16, xlab="Mean gene expression (log2+1)", ylab="Mean variance of gene expression (log2+1)", main="Variance of gene expression\nacross all samples", col=grDevices::rgb(0,100,0,50,maxColorValue=255))
  abline(0, 1)
  abline(lm(log2(dis[,2]+1)~log2(dis[,1]+1)), lwd=2, col="red")
  print(cor.test(log2(dis[,1]+1), log2(dis[,2]+1)))

  return(dis)
}

##Principal Component  Analysis and Plot functions
library(pcaGoPromoter)
pca<- function(expmatrix, groups, PCs = c(1,2), main = "PCA plot"){

  #PCA
  pcaOutput <- pcaGoPromoter::pca(expmatrix, printDropped = FALSE, scale=TRUE, center=TRUE)

  #Plot
  pcaGoPromoter::plot.pca(pcaOutput, groups, PCs = PCs, printNames = TRUE, symbolColors = TRUE, plotCI = TRUE, main=main)
}
```

***##Volcano plot***

res <- read.table("Supp-data-file-1_Back-genes-expr_V3.txt", sep = '\t', header=TRUE)

head(res)

library(calibrate)

with(res, plot(log2FoldChange, minus.log10.padj, pch=20, main="Volcano plot", xlim=c(-7,7), ylim=c(0, 20)))

with(subset(res, padj<.05 ), points(log2FoldChange, minus.log10.padj, pch=20, col="blue"))

with(subset(res, abs(log2FoldChange)>1.25), points(log2FoldChange, minus.log10.padj, pch=20, col="magenta"))

with(subset(res, padj<.05 & abs(log2FoldChange)>1.25), points(log2FoldChange, minus.log10.padj, pch=20, col="green"))

with(subset(res, padj<.01 & abs(log2FoldChange)>3), textxy(log2FoldChange, minus.log10.padj, labs=Gene, cex=.5))

with(subset(res, padj<.01 & abs(log2FoldChange)>2 & abs(log2FoldChange)<3), textxy(log2FoldChange, minus.log10.padj, labs=Gene, cex=.5))

#### STRING Network settings

Unconnected STRING network (v11) nodes were excluded, and nodes were colored according to their grouped protein function. Protein function was determined by association with KEGG Pathways (*Danio rerio*, dre04510), Reactome Pathways (*Danio rerio*, DRE-1474290, DRE-3000178, DRE-5358351, DRE-419037, DRE-5173105, DRE-381426, DRE-1592389), or PFAM Protein Domains (PF00090, PF07645) database inputs and these were selected to visualize clusters based on their assigned biological function. Clusters were manually untangled and moved for visual purposes and kept as similar as possible.

#### Gene contig, protein sequence alignment and literature look-up

ENSDARG00000068621 (*si:ch211-181d7.3*) and ENSDARG00000088274 (*si:ch211-181d7.1*) genes were searched in Ensembl (release 100) and an ortholog search was performed to determine if mammalian orthologues exist. Protein alignment and domain look-up was performed on Clustal Omega (v1.2.4)(Sievers, Wilm et al., 2011) and UniProt (v Aug-2020; accession numbers R4GE51 and E7FA88)(The UniProt, 2018) web interfaces respectively.

Gene set enrichment analysis involving human monogenic and polygenic skeletal traits and disease

##### **Orthologue mapping**

Ensembl IDs and gene symbols from differentially expressed (**supplemental data file 2**), background (**supplemental data file 1**), and whole genome (Ensembl release 99) genes lists were uploaded to dbOrtho (BioDBnet) ortholog converter (<https://biodbnet-abcc.ncifcrf.gov/db/dbOrtho.php>) using *Danio rerio* (GRCZ11) as input (‘Gene Symbol’ or ‘Ensembl ID’) and *Homo sapiens* as output (‘Ensembl ID’)(Mudunuri, Che et al., 2009). Only Ensembl IDs generating 1:1 results were kept, and duplicates of zebrafish paralogs (2:1 conversion) were removed leading to a list of single human Ensembl IDs.

Zebrafish orthologues that were successfully converted to human orthologues were subjected to exclusion criteria that addresses the existing evolutionary genomic complexity between humans and zebrafish due to an additional teleost genome duplication event leaving a subset (~20-25%) of genes with extra paralogs (HoweClark et al., 2013, Meyer & Schartl, 1999). Hence, only zebrafish genes that mapped one-to-one (e.g. sp7 to SP7) or two-to-one (e.g. ncam1a and ncam1b to NCAM1, in this case duplet Ensemble IDs were removed to have a unique human Ensembl ID) to the human genome.

***Monogenetic disease***

In conjunction with ISDS Nosology and Classification of Skeletal Disorders database, evidence of enrichment for human monogenic skeletal disease causing genes was examined using the Fisher's Exact Test of Independence under the hypergeometric distribution, using the following parameters: (A) Successes in sample - the number of zebrafish genes that were differentially expressed and that map to unique human orthologues and cause one or more skeletal disorders (either total or within each individual nosology-defined skeletal disorder groups). (B) Sample size - the number of differentially expressed zebrafish genes within the population of genes that were expressed in ontogenetic and/or regenerating genes that map to human orthologues. (C) Successes in population - the number of zebrafish genes that were expressed in ontogenetic and/or regenerating scales and that map to unique human orthologues and cause one or more skeletal disorders (either total or within each individual nosology-defined skeletal disorder groups). (D) Population - the number of zebrafish genes that are expressed in ontogenetic and/or regenerating scales and that map to human orthologue. Estimates of enrichment were obtained using the online Hypergeometric p-value calculator (<https://systems.crump.ucla.edu/hypergeometric/>).

***Polygenetic traits and disease***

MAGMA competitive gene-set analysis (de Leeuw, Mooij et al., 2015) was used to investigate whether human orthologues of zebrafish DEGs set were on average more strongly associated with polygenetic traits/disease (adult height, quantitative ultrasound derived heel bone mineral density (eBMD) and any form of self-reported, or hospital-diagnosed osteoarthritis), than all other protein coding genes in the human genome.

##### Datasets used for analysis:

Analyses of height and eBMD were performed on a sample of 378,484 unrelated white European subjects (54% female, plink1.9 / 2 (Chang, Chow et al., 2015) pairwise kinship co-efficient < 0.044) from the UK Biobank Study (Sudlow, Gallacher et al., 2015) that had valid quantitative eBMD and adult stature (i.e. height) and high-quality genome-wide HRC and 1000G/UK10K imputed data from the January 2018 release [(i.e. ~17M genetic variants with an information quality score > 0.9, minor allele frequency > 0.05%, minor allele count > 1, genotyping hard call rate > 0.95, and weak evidence of deviation from Hardy-Weinberg equilibrium (P > 1x10^-6^)]. Analyses of individuals with OA were performed on summary results statistics from a recent UK Biobank and arcoGEN GWAS meta-analysis of hospital diagnosed or self-reported OA at any skeletal site: 77,052 cases & 378,169 controls. Details of how eBMD and OA was defined and criteria for including individuals in each cohort are detailed in the original publications (Kemp, Morris et al., 2017, Morris, Kemp et al., 2018).

##### Gene-based tests of association

Gene-based tests of association were conducted in MAGMA (v1.08, https://ctg.cncr.nl/software/magma) using imputed individual level genotype data for the analyses involving adult height and eBMD, and GWAS meta-analysis summary results statistics for OA (Kindly made available by ACROGEN consortium). Analyses involving height and eBMD were further adjusted for age, sex, genotyping array, and ancestry informative principal components 1 – 20. Gene-based tests of association encompassed a multi-model approach in which the association results from different gene analysis models were combined to produce an aggregate p-value corresponding to the strength of evidence of association between each protein coding gene (+2kb upstream / 1kb downstream) and eBMD, height, or OA. The three association models included: a principal components regression model, a SNP-wise mean χ2 model [*i.e*. test statistic derived as the sum of −log(SNP p-value) for all SNPs that intersect the gene region of interest], and SNP-wise top χ2 model [(test statistic derived as the sum of −log(SNP p-value) for top SNP in the region of interest)]. The aggregate approach was chosen as it yields a more even distribution of statistical power and sensitivity over a wider range of different genetic architectures. Note: Principal components regression model could not be run for OA as the method requires individual level genotyping data that was not available. Importantly, due to incomplete genotyping coverage (i.e. no genetic variants were present in some gene), gene level associations could only be estimated for 459 / 483 DEGs, and 8,365/8,751 genes with background expression and 12,025/12,667 all mappable zebrafish – human orthologues.

##### Gene set analysis

Competitive gene set analysis was used to determine whether human orthologues of the 451 zebrafish DEG set was on average more strongly associated with adult height, eBMD or OA than all other human protein coding genes. The analysis accounted for several confounding factors including: gene size, gene density (i.e. representing the relative level of LD between SNPs in the gene) and the inverse of the mean minor allele count in the gene (i.e. to correct for potential power loss in very low minor allele count SNPs), as well the log value of these three factors. Because enrichment of the set of zebrafish DEGs was estimated relative to all human protein coding genes, we performed additional sensitivity analysis to ensure that our results were not biased by including human genes that (i) could not be confidently mapped between zebrafish and humans and (ii) that were not able to be assessed by RNA Seq as they did not pass QC (i.e.. these were not expressed in ontogenetic or regenerating scales). To achieve this, we generated two additional gene sets, one containing all mapped genes between zebrafish and humans (irrespective of whether they were expressed), and another containing the set of all genes that were mapped and expressed in ontogenetic and/or regenerating scales (i.e. background). We repeated the analysis, conditioning on both gene sets and thereby adjusted our baseline analysis for any “mapped genes vs all human gene effects” and / or “background genes vs all human gene effects”.

##### Post hoc permutation analysis

Post hoc permutation analysis was performed using supplied R-scripts with default settings. A detailed description of the methods, in addition to interpretation of resulting plots is provided in the supplementary note from de Leeuw et al 2018 (de Leeuw, Stringer et al., 2018). Briefly, post-hoc permutation analysis involved generating quantile-quantile (QQ) plots that contrasted residual Z-scores (derived from gene-based tests of association above) of each human orthologue against its expected Z-score based on all of the genes in the set, using sample quantiles of the residual Z-scores. A one sided upper 95% confidence band was generated to monitor the degree to which the Z-score for each orthologue was likely to deviate from its expected Z-score using 10,000 permutations. Orthologues that exceeded the 95% confidence band were coloured in red and all other orthologues were coloured in grey. Plots were interpreted as follows: deviation from the plot diagonal (i.e. expectation), suggested that the distribution of the observed values was different from the expected distribution. In cases where a gene set was not enriched, its residual Z-score distribution would match that of the data, and would tend to follow the plot diagonal, subject to some random variation. In contrast, if the gene set was enriched (and the association was attributable to phenotypic associations between all genes in the gene-set), the distribution of its residual Z-scores would differ from that of the data as a whole, as for every gene in the set. Consequently, the resulting distribution would deviate from the diagonal from the first gene onward. In the cases where enrichment is not due to all genes in the gene set, but rather a nested subset of genes, enrichment of the gene set would not be attributable to phenotype associations for all genes, and the resulting Z-score distribution would only differ from expectation for a subset of genes. Z-scores for the remaining genes would tend to be lower not deviate from expectation. The resulting mixture of distributions would result in a QQ-plot that, starting from the plot origin, would stay close to the diagonal initially, and deviate upwards later. The relative size of the overlap of genes in the gene set, with genes from the underlying pathway(s) that is enriched, would determine the deflection point, with smaller overlap resulting in a later (and often more pronounced) deflection.

#### **Quantitative Real-Time PCR extended methods and primers**

The cDNA qRT-PCR amplification reactions conditions were as follows: 3 min 95°C, 40 cycles of 15s 95°C and 1 min 60°C), ran on a CFX 96 (BioRad, Hercules, USA) qPCR machine, followed by a standard melting curve. Only amplicons passing the standard melting curve analysis were used for downstream calculations. The two reference genes (*eef1a1l1* and *rpl13*) and target amplicon primer sequences that have been used in this study can be found in the table below.

Primer sequences used in qPCR (supplemental table)

| Gene | Source | Forward primer (5’→3’) | Reverse primer (5’→3’) |
| --- | --- | --- | --- |
| *acp5b* | NM_001002452 | CGTCCACTGACCACAGGAAGA | AAGGATCCTGACGTCTGATTGA |
| *ctsk* | NM_001017778.1 | CTATAAAGAGATTCCTCAGGGTAACGA | ACACGGGTCCCACATTGG |
| *bgna* | NM_001002227.1 | CCAACCTCTATGCCCTGGTC | TGGGCATGGATGTCAGAAGG |
| *cgref1* | XM_017351827.2 | CGGCTTGTGTTTGTGCTCAT | CCCTTGTGTTGAGCTCTGGA |
| *col10a1a* | NM_001083827.1 | GAAATGGGGGTTGGGGTTCA | TAAACCAACTCCAGGCGCTC |
| *col11a2* | NM_001079992.2 | ATTGAAGAACAAGTCAAACCAAACC | GGCATGATCAGGAGTCCCAG |
| *col1a2* | AJ318213 | GGAAACCTGAAGAAGGCTGTGT | TGAAAGTGAAGCGGCTGTTG |
| *cx43* | NM_131038.1 | CACGCCGAAGGAACTGTCTA | GTCGAAGGCTTGTGCATGTG |
| *eef1a1l1* | NM_131263.1 | CTGGAGGCCAGCTCAAACAT | TCAAGAAGAGTAGTACCGCTAGCATTAC |
| *entpd5a* | XM_679770.8 | GCGGAACCCCAGATGGTT | CACCTGCAGCACCTCTTGGT |
| *ihha* | NM_001034993.2 | GCTTGTGTACAGCGAGGTCA | CAAGCTCAGTTTCGCACCAG |
| *mmp9* | NM_213123.1 | AAATCTGTGTTCGTGACGTTTCCT | GCCGTAACGCTTCAGATACTCAT |
| *prdm5* | NM_001002301.2 | GACGTGTGTGACCTGTCCTT | TCGGCCTGTTGGGATTATGG |
| *rpl13* | NM_212784.1 | TCTGGAGGACTGTAAGAGGTATGC | AGACGCACAATCTTGAGAGCAG |
| *smpd3* | NM_001122750.1 | TTTCGAAGGGTCACCTCGTG | GTGAAGAGCGAATGCTGCTG |
| *spp1* | NM_001002308.1 | GGCGGCTTGACATTTGTGAG | GTCCCCAACGGGAACAATCT |
| *sp7* | NM_212863.2 | GGATACGCCGCTGGGTCTA | TCCTGACAATTCGGGCAATC |
| *tmem97* | NM_001326398.1 | TCTTCGCGTCCTGGAAATCAT | CCTCGCAGAACACAAACGAC |
