## Supplementary material for "The transcriptome of regenerating zebrafish scales identifies genes involved in human bone disease": supplemental tables and figures.docx (from another version of this preprint: version 1, 2020-10-08)

**Supplemental tables and figures, Bergen et al.**

| Cluster # | *Danio rerio* gene symbol | *Danio rerio* Ensembl ID | Log_2_ Fold Change | False Discovery Rate | Wikigene description |
| --- | --- | --- | --- | --- | --- |
| 1 | *si:ch211-204c21.1*  (*LOC553492*) | ENSDARG00000053091 | 1.564 | 4.26E-04 | B double prime 1, subunit of RNA polymerase III transcription initiation factor IIIB (2342 aa) |
| 1 | *myo6b* | ENSDARG00000042141 | 1.382 | 1.58E-03 | myosin VIb |
| 1 | *sh3gl3b* | ENSDARG00000007302 | 1.415 | 2.04E-02 | SH3-domain GRB2-like 3b |
| 1 | *dnm1b* | ENSDARG00000009281 | 1.672 | 2.19E-02 | dynamin 1b |
| 2 | *rhoua* | ENSDARG00000019709 | 1.821 | 5.73E-11 | ras homolog family member Ua |
| 2 | *bcr* | ENSDARG00000079286 | 1.663 | 2.65E-10 | BCR activator of RhoGEF and GTPase |
| 2 | *gdi1* | ENSDARG00000056122 | 1.381 | 1.11E-05 | GDP dissociation inhibitor 1 |
| 2 | *arhgef9a* | ENSDARG00000061746 | 1.943 | 6.51E-05 | Cdc42 guanine nucleotide exchange factor (GEF) 9a |
| 2 | *srgap1b* | ENSDARG00000045789 | 2.050 | 3.74E-04 | SLIT-ROBO Rho GTPase activating protein 1b |
| 2 | *arhgef40* | ENSDARG00000078979 | 1.387 | 1.36E-03 | Rho guanine nucleotide exchange factor (GEF) 40 |
| 3 | *mvda* | ENSDARG00000099336 | -1.836 | 2.51E-05 | mevalonate (diphospho) decarboxylase a |
| 3 | *mvk* | ENSDARG00000004130 | -1.920 | 8.04E-05 | mevalonate kinase |
| 3 | *hmgcs1* | ENSDARG00000103025 | -1.831 | 0.00137 | 3-hydroxy-3-methylglutaryl-CoA synthase 1 (soluble) |
| 3 | *tm7sf2* | ENSDARG00000032816 | -1.913 | 0.000689 | transmembrane 7 superfamily member 2 |
| 3 | *sigmar1* | ENSDARG00000011418 | -1.500 | 0.00115 | sigma non-opioid intracellular receptor 1 |
| 3 | *cyp46a1.3* | ENSDARG00000089177 | 2.085 | 0.000546 | cytochrome P450, family 46, subfamily A, polypeptide 1, tandem duplicate 3 (zgc:136808) |
| 3 | *dhcr7* | ENSDARG00000103226 | -2.137 | 0.000183 | 7-dehydrocholesterol reductase |
| 3 | *cyp2r1* | ENSDARG00000056587 | 2.380 | 1.35E-08 | cytochrome P450, family 2, subfamily R, polypeptide 1 |
| 3 | *cyp24a1* | ENSDARG00000103277 | -2.182 | 0.000212 | cytochrome P450, family 24, subfamily A, polypeptide 1 |
| 4 | *mybpc2b* | ENSDARG00000021265 | 6.520 | 1.04E-18 | myosin binding protein C, fast type b |
| 4 | *tmod4* | ENSDARG00000020890 | 1.883 | 3.91E-05 | tropomodulin 4 (muscle) |
| 4 | *tpm4a* | ENSDARG00000023963 | 1.276 | 9.81E-05 | tropomyosin 4a |
| 4 | *tpm4b* | ENSDARG00000019128 | 1.581 | 1.88E-04 | tropomyosin 4b |
| 4 | *tpm1* | ENSDARG00000087402 | 1.408 | 3.56E-04 | tropomyosin 1 (alpha) |
| 4 | *fibina* | ENSDARG00000039949 | 1.510 | 1.34E-02 | fin bud initiation factor a |
| 4 | *myl10* | ENSDARG00000062592 | 3.180 | 1.71E-02 | myosin, light chain 10, regulatory |
| 4 | *myl6* | ENSDARG00000115496 | 1.508 | 1.25E-02 | myosin, light chain 6, alkali, smooth muscle and non-muscle |
| 4 | *mylk3* | ENSDARG00000076348 | 2.320 | 2.92E-03 | myosin light chain kinase 3 |
| n.b. | *hmgcra* (cluster 3) has been omitted as it was not a DEG and was added by STRING as a ‘high confidence predicted functional partner’. | | | | |

**Table S1: DEGs identified belonging to isolated clusters (1-4) with no identified gene ontology enrichment in the STRING analysis.**

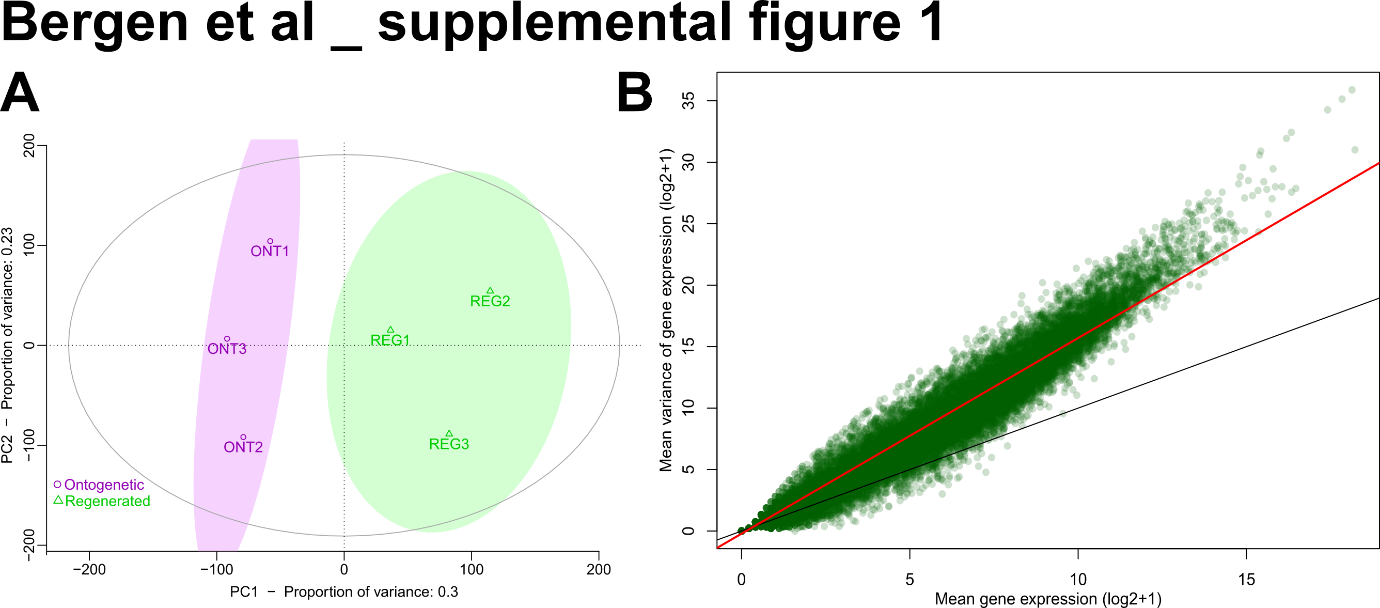

**Figure S1: Ontogenetic and regenerating differential expression values are clustered together. A)** Principle component analysis (PCA) shows that ontogenetic and regenerating samples show similar variance in their respective groups. **B)** Linear variance analysis showed high level of correlation, Pearson’s correlation, r^2^=0.9714.

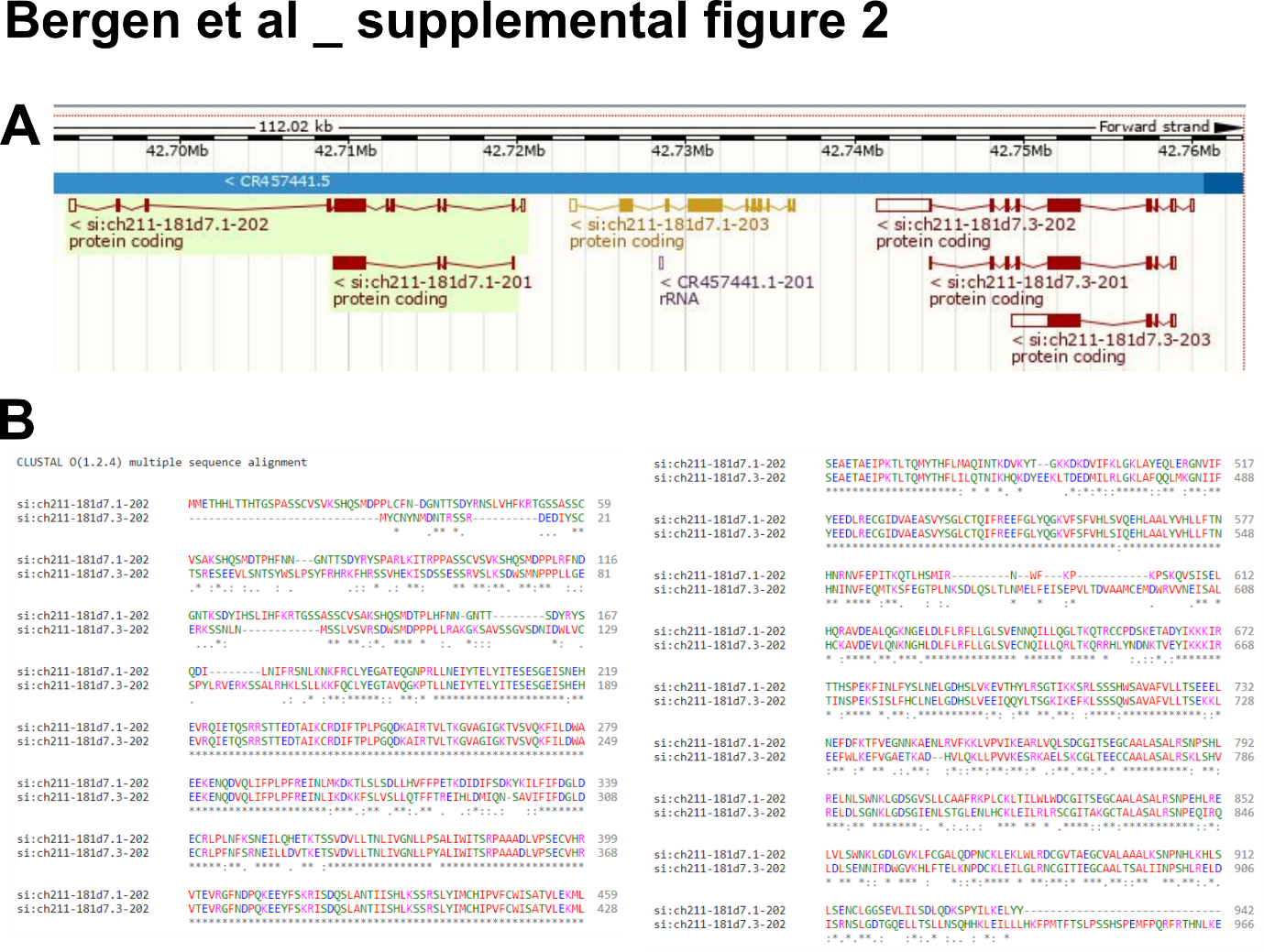

**Figure S2: Two top down regulated genes are on the same genomic contig but are not the same protein. A)** Screenshot of Ensembl release 100 website showing *si:ch211-181d7.1* and *si:ch211-181d7.3* protein coding genes are on the same genomic contig (CR457441.5) located on chromosome 15. **B)** Clustal Omega protein sequence alignment showing only high sequence similarity ranging from amino acid ~200 to ~580 (*si:ch211-181d7.1*) between the two proteins.

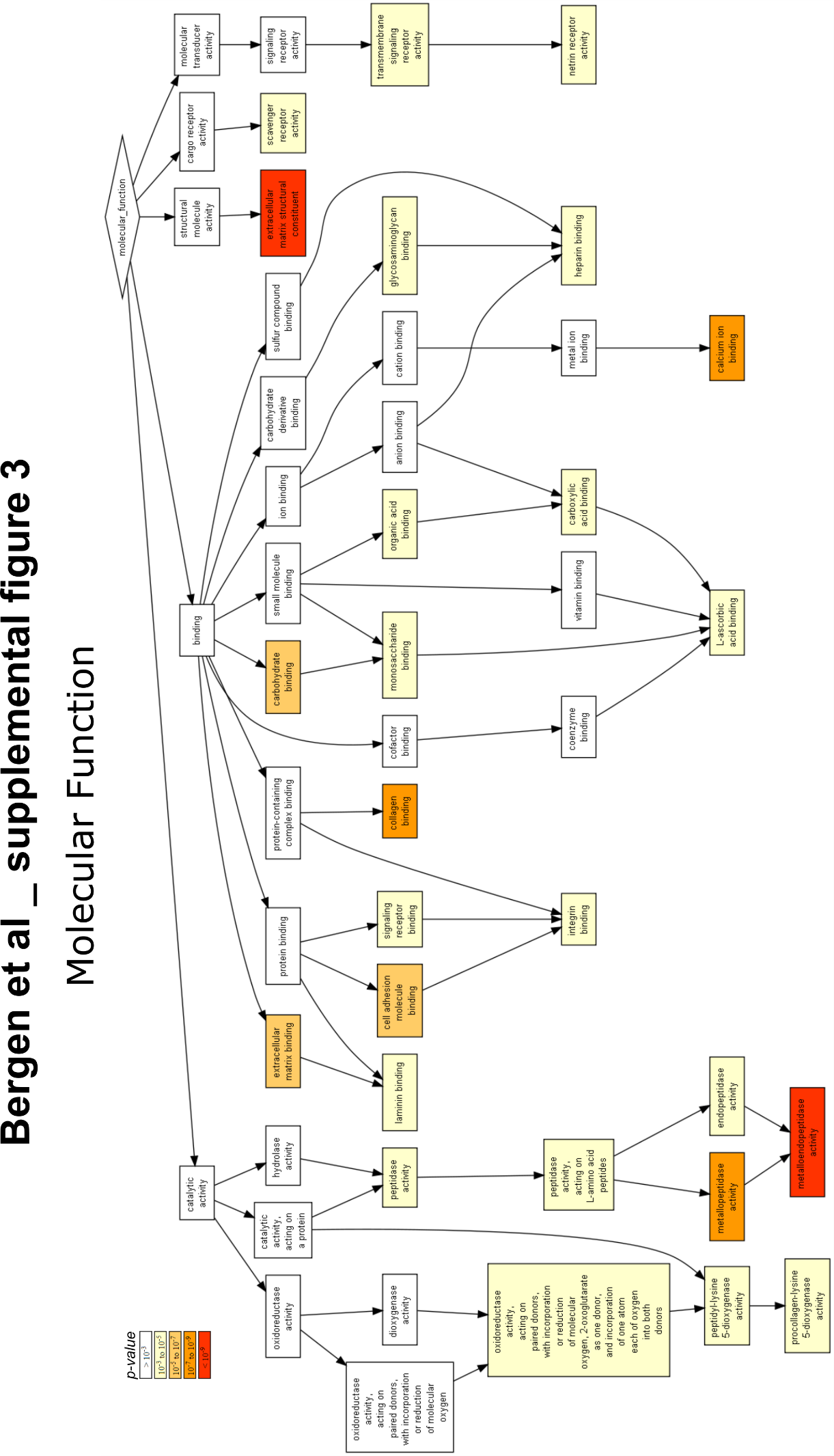

**Figure S3: Hierarchical clustering of ‘Molecular Function’ enriched gene ontology terms.** Exported from GOrilla web interface showing GO enrichment of DEGs against background expression dataset.

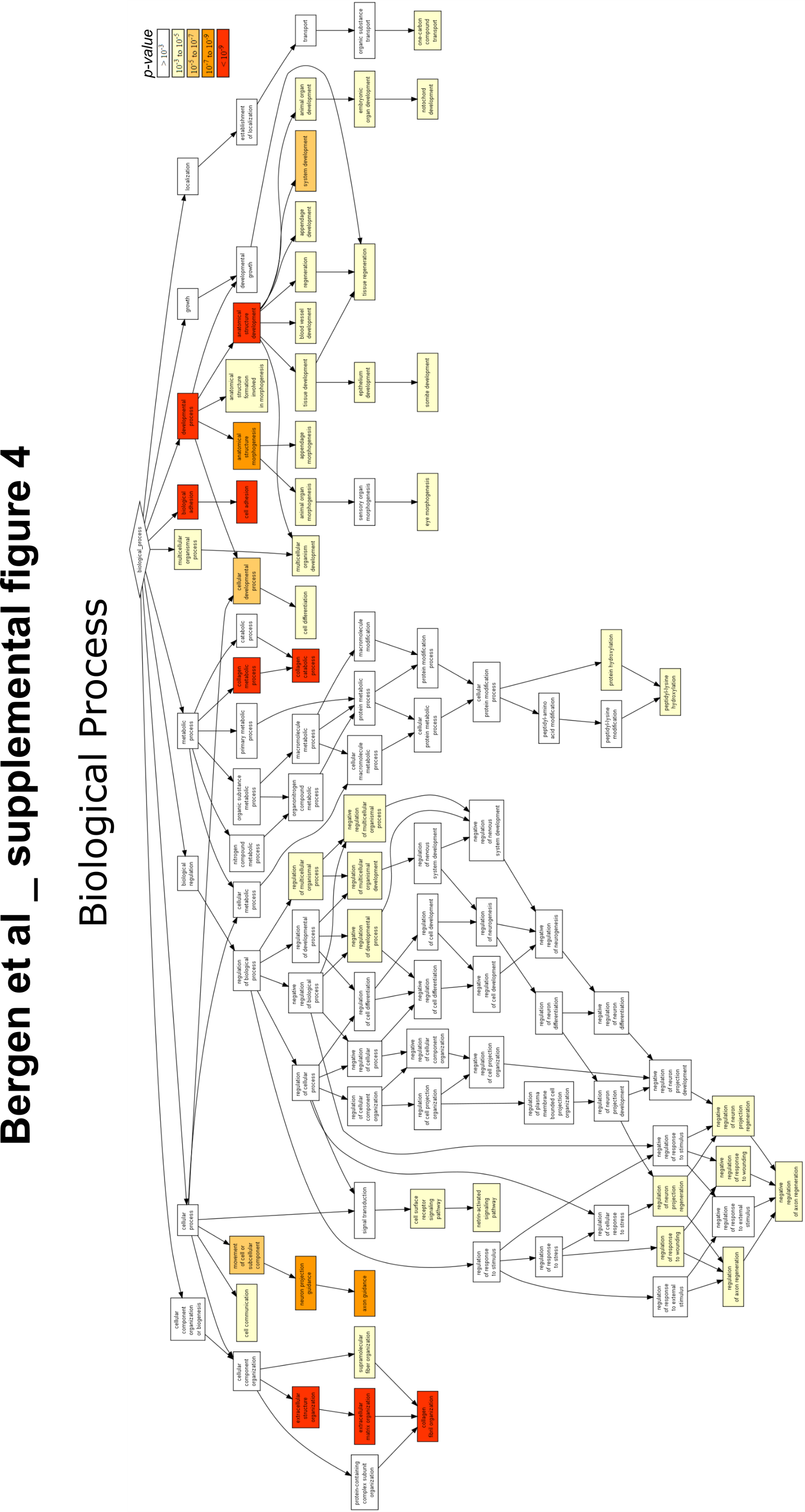

**Figure S4: Hierarchical clustering of ‘Biological Process’ enriched gene ontology terms.** Image was exported from the GOrilla web interface showing GO enrichment of DEGs against background expression dataset.

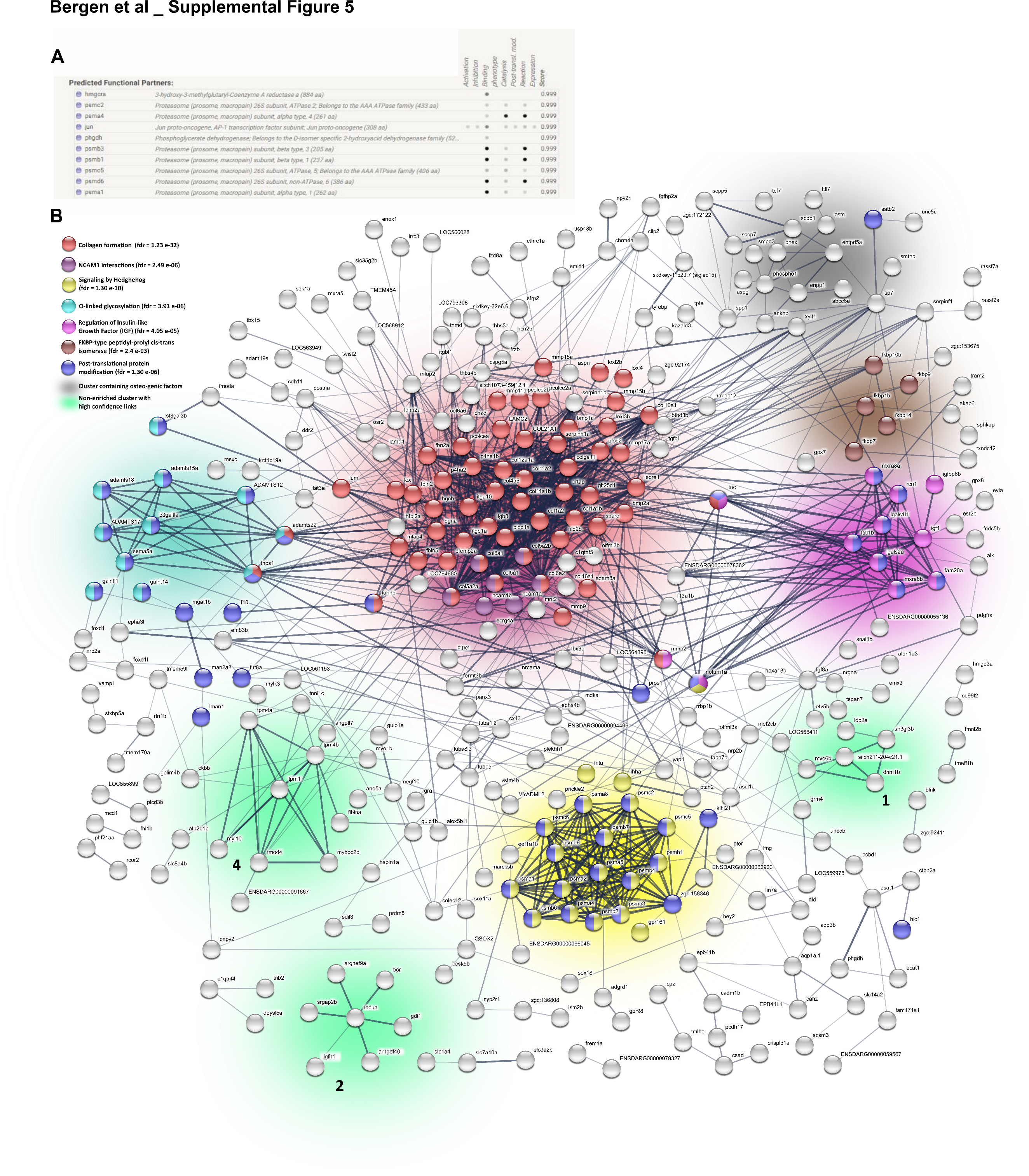

**Figure S5: STRING Network analysis showing high protein-protein interaction connectivity of DEGs. A)** Proteins added to the network that showed high interaction scores (=0.999) with DEGs that were added to the ‘high confidence’ (>0.7) and ‘medium confidence’ (>0.4) protein-protein interaction networks (PPI). **B)** ‘Medium confidence’ (>0.4) STRING Network of DEGs showing high connectivity between networks. Note that only cluster 2 is not connected to the overall network. PPI enrichment P-value for this network: < 1.0e-16 with a 2.8-fold enrichment in number of connections between nodes.

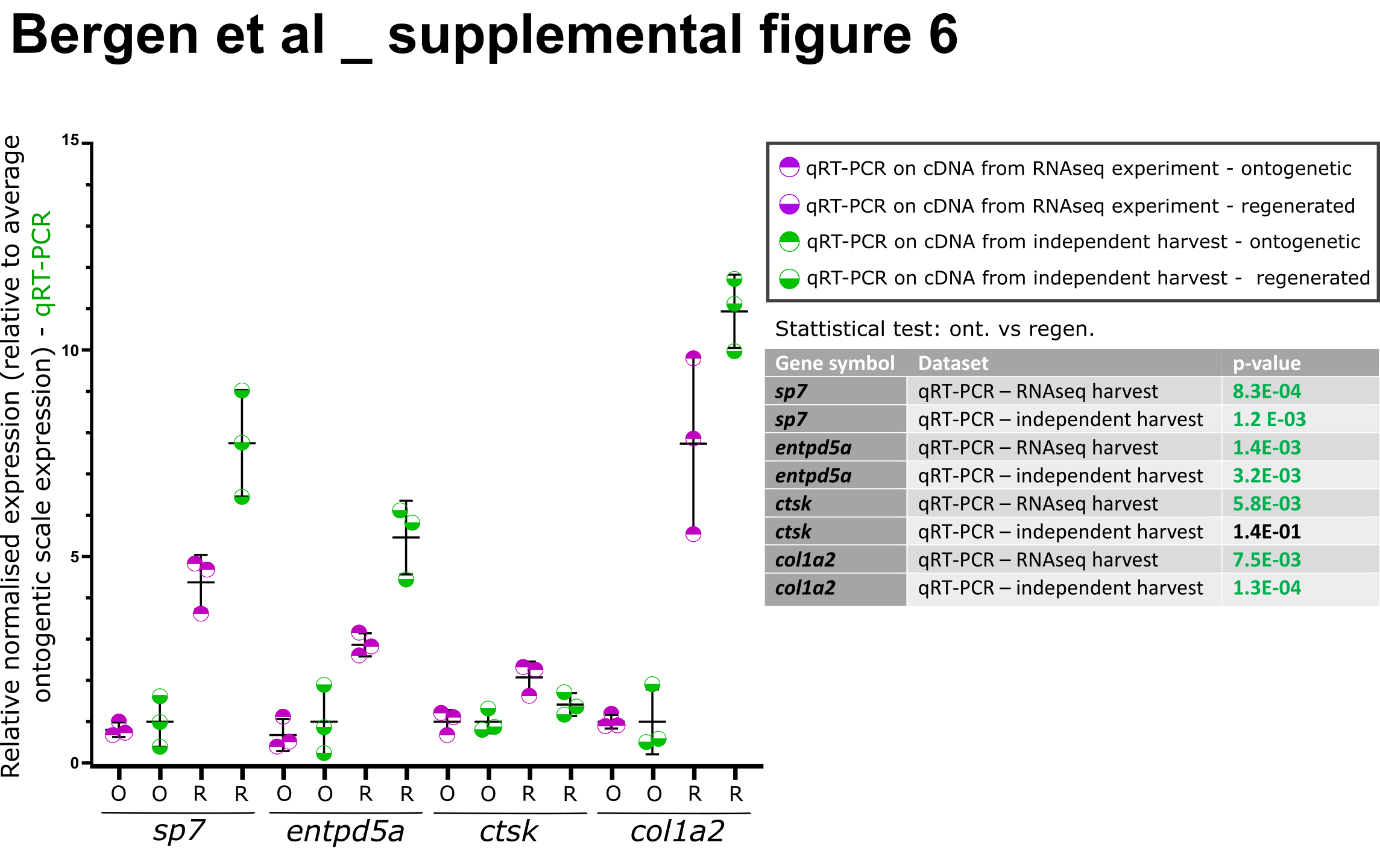

**Figure S6: Quantitative Real-Time PCR analysis of RNA expression of bone markers between RNA isolated from the RNA-sequencing scale harvest and an additional independent harvest.** Expression profiles show similar expression patterns of the selected amplicons between the two RNA isolations from ontogenetic and regenerating scales.

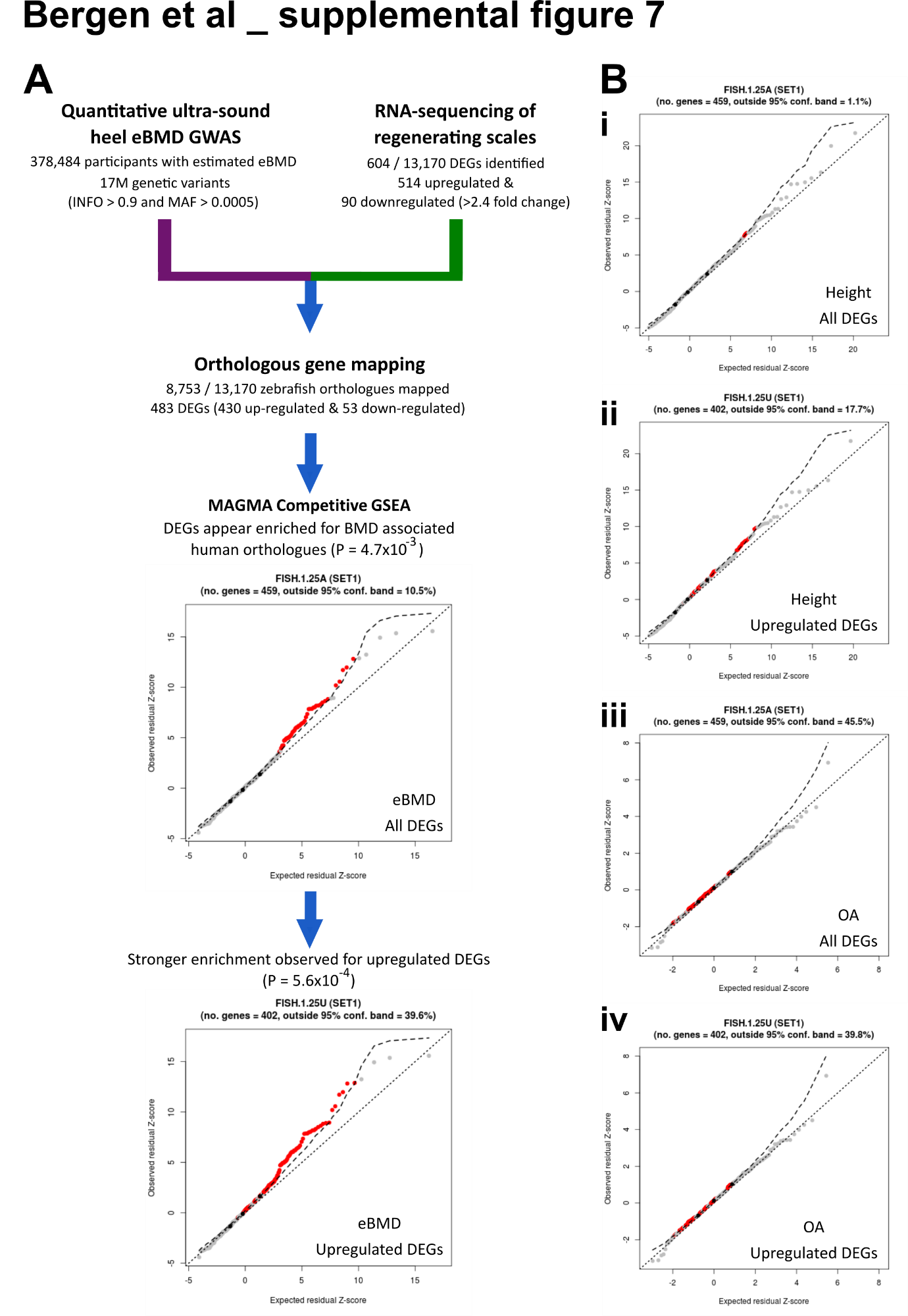

**Figure S7: MAGMA competitive gene set enrichment analysis involving human polygenetic traits and disease. (A)** MAGMA competitive gene set enrichment analysis was used to investigate the relationship between genetic variation surrounding mappable human orthologs of 459 differentially expressed zebrafish genes and quantitative ultrasound derived heel bone mineral density (eBMD) measured in 378,484 unrelated white European adults from the UK Biobank Study. Strong evidence of enrichment was observed, suggesting that human orthologues of zebrafish DEGs were on average more strongly associated with the eBMD than all other human protein coding genes in the genome. Post-hoc permutation analysis involved generating quantile-quantile (QQ) plots that contrasted residual Z-scores of differentially expressed orthologue (derived from gene-based tests of association) against its expected Z-score based on the quantile of all orthologues, with the 25th, 50th and 75th percentile denoted by black circles. A one sided upper 95% confidence band was generated to monitor the degree to which the Z-score for each orthologue was likely to deviate from its expected Z-score (denoted by the black dashed line). Orthologues that exceeded this confidence band were coloured in red and all other orthologues were coloured in grey. The proportion of orthologues exceeding the 95% confidence band was reported. Post hoc analysis revealed a mixture of distributions that resulted in a QQ-plot that, (starting from the plot origin) stayed close expectation (denoted by the solid black diagonal line) for most of the differentially expressed orthologues, and deviated upwards towards the end. This pattern, characterised by the late deflection point, suggested that enrichment was not attributable to all differentially expressed orthologues, but rather that the Z-score distribution differed from expectation for a subset of orthologues. To further investigate this finding, orthologues were stratified according to whether they we up- or downregulated, and both sets were re-analysed separately. Stronger enrichment for was observed for the 402 up-regulated orthologues as compared with the set of all differentially expressed orthologues, and no evidence of enrichment was observed for the downregulated orthologues (data not shown). Subsequent post hoc analysis revealed that deviation from expectation occurred earlier and a larger proportion of orthologues exceeded the 95% confidence band (i.e. 40% vs 11% as compared with all differentially expressed orthologues). These findings suggested that human orthologues of many, but not all upregulated DEGs are likely to regulate bone homeostasis in human populations. **(B)** MAGMA gene-set enrichment analysis involving height and self-reported or diagnosed osteoarthritis suggested that DEGs were enriched for human orthologues associated with both traits. Inspection of the QQ plots for height **(i)** and osteoarthritis **(iii)** analysis suggested that enrichment was less pronounced as compared with eBMD, and that in both cases not attributable to all differentially expressed orthologues, To further investigate these findings, orthologues were stratified according to whether they we up- or downregulated, and each set was re-analysed. Enrichment for height associated orthologues was stronger for the 402 up-regulated orthologues as compared with the set of all orthologues, and no evidence of enrichment was observed for the downregulated DEGs (data not shown). Subsequent post hoc permutation analysis revealed a mixture of distributions that was slightly less pronounced **(ii)**, and the deflection point from expectation that started earlier (as compared to the analysis involving all orthologues). These findings suggested that enrichment for height associated orthologues was attributable a moderate proportion (but not all) upregulated orthologues. Analysis involving osteoarthritis revealed that the magnitude of enrichment for upregulated orthologues was similar to the analysis involving all differentially expressed orthologues. QQ plots were similar for both sets of analyses (**ii and iv**), revealing a mixture of distributions suggesting that enrichment was likely attributable to many, but not all differentially expressed orthologues.

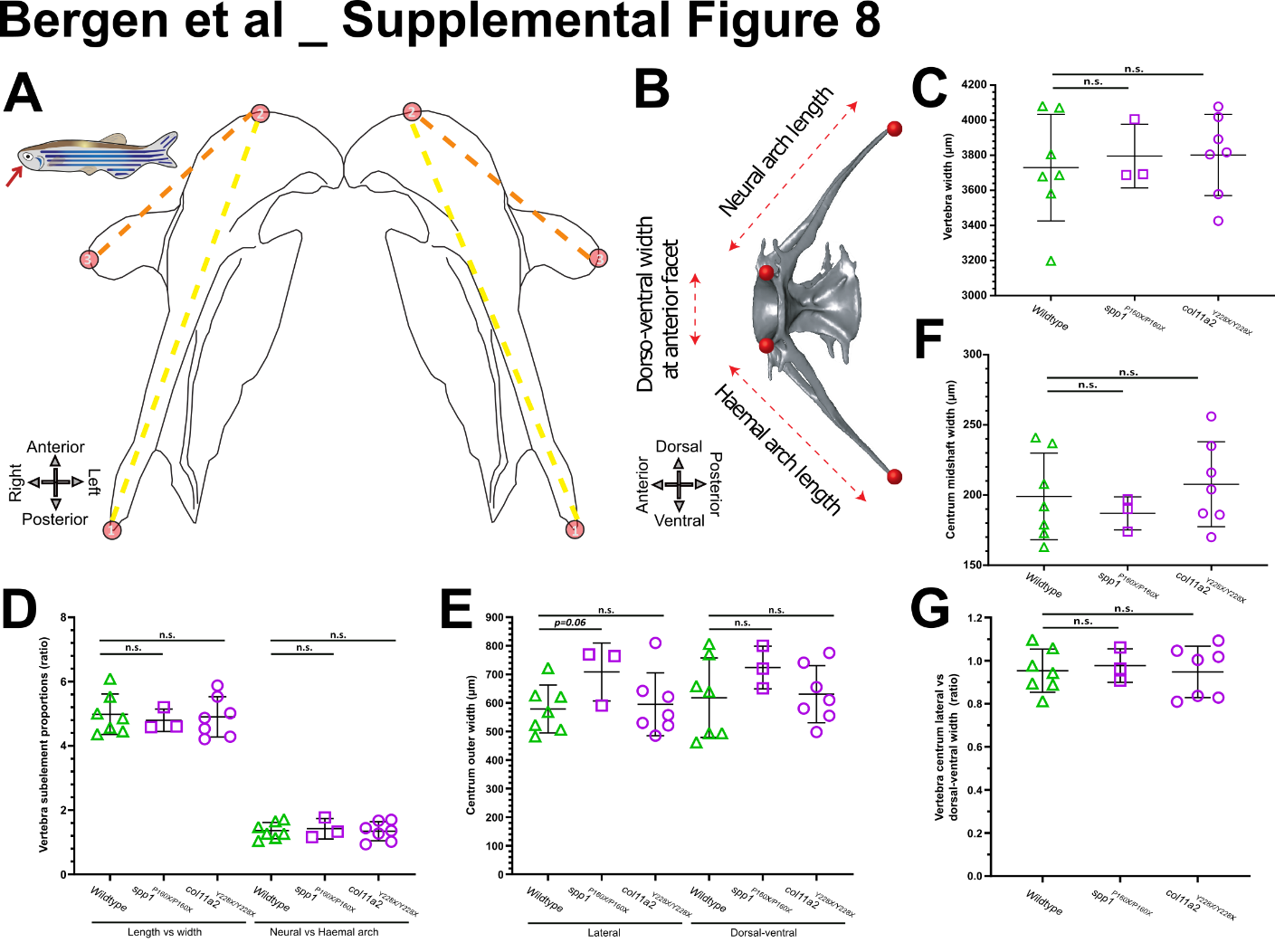

**Figure S8: Histomorphological measurements of skeletal elements from wildtype, *col11a2^Y228X/Y228X^* and *spp1^P160X/P160X^* fish. A)** Schematic drawing of the landmark points to measure histomorphological parameters of the ventral lower jaw (red arrow). Landmark point 1-2 is length, and 2-3 is width. **B)** Landmark points of a representative 3D micro-CT image from a caudal vertebra used for measuring histomorphological parameters in this study. **C-G)** Measurements of the various caudal vertebrae (number 11-13) element parameters of wildtype, *col11a2^Y228X/Y228X^* and *spp1^P160X/P160X^* fish. Mean values of length (anterior to posterior) and mean width (for whole vertebra element: haemal arch length + neural arch length + anterior facet centrum width) of each individual were used.
